# A pair of transporters controls mitochondrial Zn^2+^ levels to maintain mitochondrial homeostasis

**DOI:** 10.1101/2021.07.21.453180

**Authors:** Tengfei Ma, Liyuan Zhao, Jie Zhang, Ruofeng Tang, Xin Wang, Nan Liu, Qian Zhang, Fengyang Wang, Meijiao Li, Qian Shan, Yang Yang, Qiuyuan Yin, Limei Yang, Qiwen Gan, Chonglin Yang

## Abstract

Zn^2+^ is required for the activity of many mitochondrial proteins, which regulate mitochondrial dynamics, apoptosis and mitophagy. However, it is not understood how the proper mitochondrial Zn^2+^ level is achieved to maintain mitochondrial homeostasis. Using *Caenorhabditis elegans*, we reveal here that a pair of mitochondrion-localized transporters controls the mitochondrial level of Zn^2+^. We demonstrate that SLC-30A9/ZnT9 is a mitochondrial Zn^2+^ exporter. Loss of SLC-30A9 leads to mitochondrial Zn^2+^ accumulation, which damages mitochondria, impairs animal development and shortens the life span. We further identify SLC-25A25/SCaMC-2 as an important regulator of mitochondrial Zn^2+^ import. Loss of SLC-25A25 suppresses the abnormal mitochondrial Zn^2+^ accumulation and defective mitochondrial structure and functions caused by loss of SLC-30A9. Moreover, we reveal that the endoplasmic reticulum contains the Zn^2+^ pool from which mitochondrial Zn^2+^ is imported. These findings establish the molecular basis for controlling the correct mitochondrial Zn^2+^ levels for normal mitochondrial structure and functions.

**Summary:** Zn^2+^ is a trace ion essential for the function of many mitochondrial proteins. It is not known how mitochondrial Zn^2+^ levels are regulated. Ma at al. identify transporters that mediate mitochondrial Zn^2+^ export and import to maintain mitochondrial homeostasis.

## Introduction

Zinc, in the ion form (Zn^2+^), is an essential trace element for organisms. Zn^2+^ plays important roles in a wide range of physiological and cellular processes such as development, metabolism, DNA synthesis, and transcription (Colvin et al., 2010; Kambe et al., 2015). In humans, Zn^2+^ binds to about 10% of total proteins, acting either as a structural component of functional proteins (e.g. zinc fingers) or as a cofactor of catalytic enzymes including oxidoreductases, transferases, hydrolyases, lyases, isomerases, and ligases (Andreini et al., 2006; Colvin et al., 2010). In addition, Zn^2+^ functions as an important signaling ion (Frederickson et al., 2005; Fukada et al., 2011; Yamasaki et al., 2007). Intracellular Zn^2+^ promotes the expression and secretion of nerve growth factor (NGF) and early growth response factor 1 (EGR1), and promotes the activation of extracellular signal-regulated kinase 1/2 (ERK1/2) (Park and Koh, 1999). In synapses, synaptic Zn^2+^ permeates Ca^2+^- and Zn^2+^-permeable GluR2-lacking AMPA receptors (AMPAR_Ca-Zn_s) and VGCCs (voltage-gated Ca^2+^ channels) to activate TrkB through either Src kinase- or metalloproteinase-dependent pathways, which in turn triggers synaptic potentiation (Huang et al., 2008). In *C. elegans*, physiological concentrations of Zn^2+^ were found to modulate Ras-mediated signaling (Bruinsma et al., 2002; Yoder et al., 2004). Zn^2+^ deficiency has been found to associate with poor growth and development, congenital neurological and immune disorders, defective wound healing, chronic inflammation, alopecia, persistent diarrhea, and bleeding (Kambe et al., 2015; Mammadova-Bach and Braun, 2019), to name a few. On the other hand, excess Zn^2+^ is highly toxic. Zn^2+^ accumulation is known to potently induce neuronal cell death in ischemia and blunt head trauma, and is associated with Alzheimer’s disease (AD) (Sensi et al., 2009).

Because either insufficient or excessive Zn^2+^ is deleterious to the cell, the cellular Zn^2+^ levels need to be tightly controlled. Cellular uptake and export of Zn^2+^ are carried out by two families of Zn^2+^ transporting proteins: Zrt, irt-related proteins (ZIPs), which are responsible for Zn^2+^ influx into the cytoplasm, and Zn^2+^ transporters (Znt), which mediate Zn^2+^ efflux from the cytosol (Kambe et al., 2015). These Zn^2+^ transporters usually localize to the plasma membrane or organelle membranes. In the cytosol, about 5-15% of Zn^2+^ is sequestered by binding to metallothioneins (MTs), oxidation of which leads to release of free Zn^2+^ (Colvin et al., 2010; Kambe et al., 2015). In addition, cellular Zn^2+^ is compartmentalized in subcellular organelles, including the nucleus, endoplasmic reticulum (ER), Golgi, endosomes and lysosomes (Colvin et al., 2010). Shuttling of Zn^2+^ between these intracellular organelles and the cytosol is also mediated by Znt/ZIP transporters (Kambe et al., 2015). Surprisingly, it is not known to date how Zn^2+^ is transported into and out of mitochondria, the centers of cellular metabolism. Zn^2+^ is required for the activity of a wide variety of mitochondrial proteins, including OMA1 and YME1L, the enzymes responsible for cleavage of OPA1, which mediates the fusion of mitochondrial inner membranes (Head et al., 2009). It was found that ROS-induced lysosomal Zn^2+^ release and mitochondrial Zn^2+^ increase trigger mitochondrial division by promoting mitochondrial recruitment of Drp1 (Abuarab et al., 2017). A possible Drp1-ZIP1 interaction likely triggers the entry of Zn^2+^ into mitochondria, which reduces the mitochondrial membrane potential (MMP) and induces mitochondrial division or mitophagy (Cho et al., 2019). Mitochondria are known to play an important role in Zn^2+^ homeostasis. Mitochondrial uptake of Zn^2+^ promotes the clearance of cytosolic Zn^2+^ in neurons undergoing excitotoxicity (Dineley et al., 2005; Paoletti et al., 1997; Sensi et al., 2009). Nevertheless, abnormal accumulation of Zn^2+^ in mitochondria has multiple adverse effects by causing loss of mitochondrial membrane potential, inhibition of complexes I and III of the electron transport chain, inhibition of α-ketoglutarate dehydrogenase, and production of ROS (Dineley et al., 2003; Sensi et al., 2009). Zn^2+^ accumulation in mitochondria also activates mitochondrial permeability transition pore (MPTP), leading to release of proapoptotic factors and cell death (Bossy-Wetzel et al., 2004; Jiang et al., 2001; Malaiyandi et al., 2005a; Sensi et al., 2009). In addition, Zn^2+^ inhibits mitochondrial movement in neurons (Malaiyandi et al., 2005a). Elevated mitochondrial Zn^2+^ might also promote PINK/Parkin-mediated mitophagy in response to hypoxia-oxygenation (Bian et al., 2018).

Mitochondria are the powerhouses of the cell and coordinate the tricarboxylic acid (TCA) cycle, oxidative phosphorylation (OXPHOS), and ATP production to provide energy for cell survival and functions. At the same time, mitochondria are central to cell death, differentiation, and the innate immune response (Spinelli and Haigis, 2018). Mitochondria undergo many homeostatic processes, such as fusion, fission, mitophagy and the mitochondrial unfolded protein responses (UPR^mt^), to maintain their structure and functions (Mishra and Chan, 2016; Pickles et al., 2018; Youle and van der Bliek, 2012). Despite the fact that many mitochondrial proteins require the presence of Zn^2+^ for their functions, it is largely unknown how mitochondrial structure and functions are affected when mitochondrial Zn^2+^ homeostasis is disrupted. Particularly, the identities of mitochondrion-localized Zn^2+^ transporters remain to be revealed. In this study, we use *C. elegans* as a model to demonstrate that SLC-30A9/Znt-9 is the mitochondrial Zn^2+^ exporter, loss of which causes mitochondrial Zn^2+^ accumulation and disruption of mitochondrial structure and functions. We further identify the Ca^2+^-activated Mg^2+^-ATP transporter SLC-25A25 as an important regulator of mitochondrial Zn^2+^ import, loss of which suppresses the accumulation of mitochondrial Zn^2+^. Moreover, we reveal that the ER serves as the Zn^2+^ reservoir, from which mitochondrial Zn^2+^ is imported. Our findings thus provide molecular insights into the mechanisms that control mitochondrial Zn^2+^ levels for maintenance of mitochondrial homeostasis.

## Results

### Loss of *slc-30A9* leads to mitochondrial abnormality

To identify regulators that are important for mitochondrial homeostasis, we performed genetic screens to search for mutants that display abnormal mitochondrial morphology, using *C. elegans* adult animals carrying an integrated array (*yqIs157*) that expresses mitochondrion-targeted GFP (Mito-GFP) in hypodermal cells (Tang et al., 2020; Zhou et al., 2019). Among the mutants we obtained, 5 of them (*yq158*, *yq166*, *yq172*, *yq189*, *yq212*) displayed similar abnormally enlarged spherical Mito-GFP-positive structures with varying sizes in the hypodermis, in contrast to the predominantly tubular Mito-GFP-labeled structures in wild-type (N2) animals (Fig. 1 A). These mutants failed to complement one another, suggesting that they affected the same gene. We thus used the *yq158* mutant to carry out further analysis. In *yq158* animals, the abnormal spherical structures were also positive for mCherry-tagged TOMM-20 (TOMM-20::mCh, outer mitochondrial membrane) (Fig. 1 B) and GFP-fused F54A3.5 (F54A3.5::GFP, the Mic10 subunit of the MICOS complex, inner mitochondrial membrane) (Fig. S1 A). This indicates that the large spherical structures were indeed abnormal mitochondria. Using another integrated array, *hqIs181*, which expresses GFP fused with the mitochondrial localization sequence (mtLS::GFP) in multiple tissues (Zhou et al., 2019), we found that abnormal spherical mitochondria also existed in muscle and intestinal cells in *yq158* mutants, in contrast to the tubular mitochondria in these tissues in N2 animals (Fig. 1 C). To determine how mitochondrial ultrastructure was affected, we performed transmission electron microscopy (TEM) analysis. N2 animals contained tubular or small globular mitochondria in both hypodermal and muscle cells. In contrast, *yq158* mutants had greatly enlarged mitochondria with severely damaged cristae (Fig. 1 D), consistent with the observations using fluorescence markers. Under TEM, abnormal mitochondria were observed in the germline, sperm, oocytes and embryos in *yq158* mutants (Fig. S1 B). In addition, we examined the time-course of appearance of abnormal mitochondria in larval development. In N2 animals, mitochondria in the hypodermis were mostly tubular and well connected from larval stage 1 (L1) to adults. In *yq158* mutants, however, the abnormal spherical mitochondria were observed as early as in L1 animals, and became predominant in adults (Fig. S1 C). Taken together, these findings suggest that the gene affected in *yq158* and the other 4 mutants is important for mitochondrial homeostasis.

**Figure 1.**
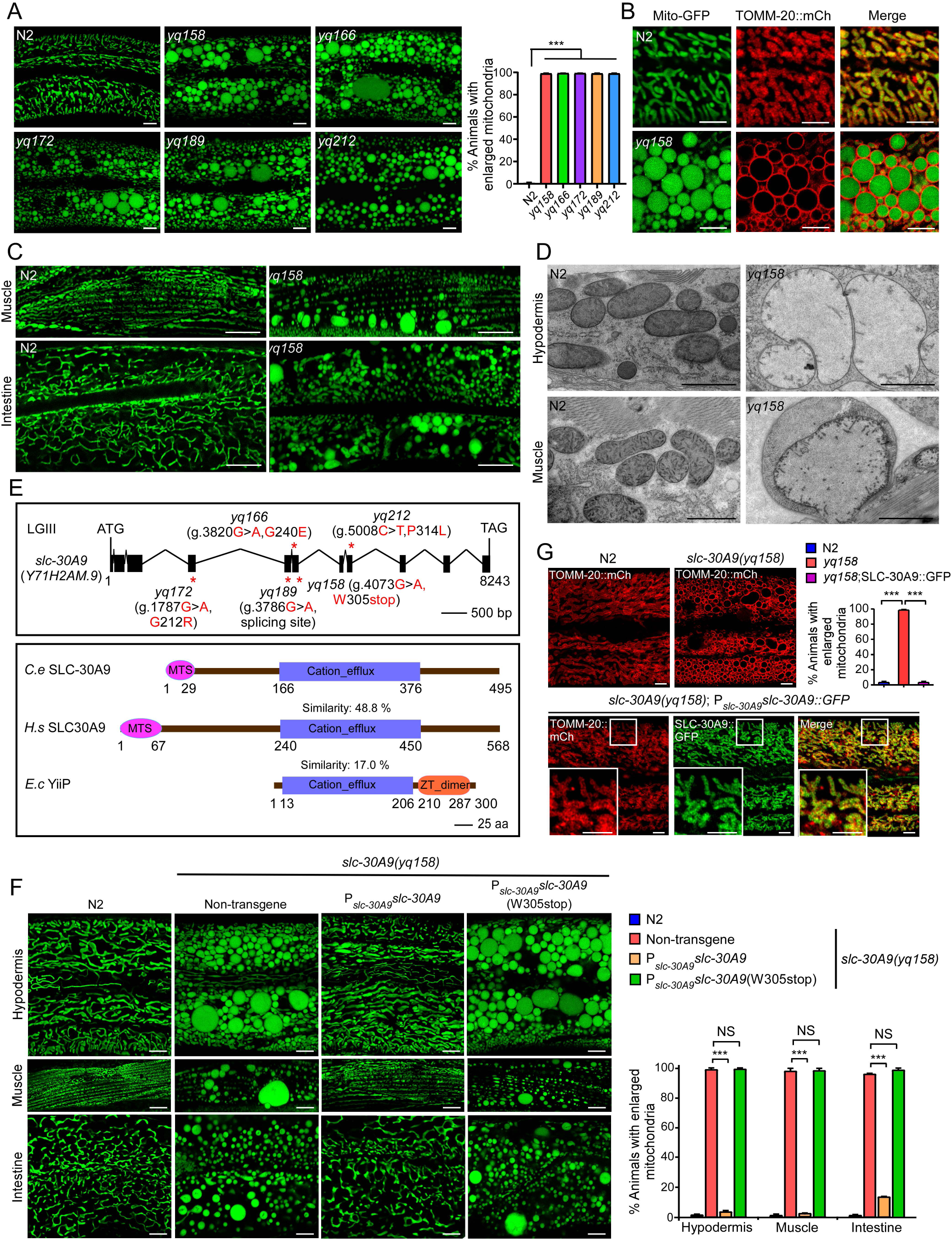
Mutations in *slc-30A9* cause abnormal mitochondrial enlargement in *C. elegans*. (**A**) Representative images (left) and quantification (right) of Mito-GFP-labeled mitochondria in the hypodermis of adult wild type (N2) and mutants (*yq158*, *yq166*, *yq172*, *yq189* and *yq212*). Mito-GFP is expressed from the chromosomally integrated array *yqIs157*. Bars, 5 μm. Abnormally enlarged mitochondria with area ≥12 μm^2^ were quantified. ≥90 animals were scored for each genotype. Comparisons are between N2 and mutants. (**B**) Images of mitochondria labeled with Mito-GFP and TOMM-20::mCh in the hypodermis of N2 and *slc-30A9*(*yq158*) animals. Bars, 5 μm. (**C**) Images of mtLS::GFP-labeled structures in muscle and intestinal cells in N2 and *slc-30A9*(*yq158*) mutant animals. mtLS::GFP is expressed from the chromosomally integrated array *hqIs181*. Bars, 5 μm. (**D**) Representative TEM images of mitochondria in the hypodermal and muscle cells in adult N2 and *slc-30A9*(*yq158*) mutant animals. Bars, 1 μm. (**E**) Top: schematic representation of the *slc-30A9* gene. Filled boxes represent exons and thin lines indicate introns. The point mutations of *slc-30A9* are indicated with asterisks. Bottom: comparison of *C. elegans* SLC-30A9 with human (*H.s*) SLC30A9 and bacterial (*E.c*) Yiip. MTS: Mitochondrial targeting sequence, ZT: Zinc transporter. (**F**) Images (left) and quantification (right) of the rescuing effects of SLC-30A9(P*_slc-30A9_slc-30A9*) and SLC-30A9(W305stop) (P*_slc-30A9_slc-30A9*(W305stop)) on *slc-30A9*(*yq158*) mitochondria in hypodermal, muscle and intestinal cells. Bars, 5 µm. ≥90 animals were scored for each transgene. (**G**) Images (top left, bottom) and quantification (top right) of the rescuing effects of SLC-30A9::GFP (P*_slc-30A9_slc-30A9*::GFP) on *slc-30A9*(*yq158*) hypodermal mitochondria labeled with TOMM-20::mCh. Boxed regions are magnified (2.5×) in the insets. ≥90 animals were scored for each genotype. Bars, 5 µm. For all quantifications, error bars represent SEM. *, P < 0.05; **, P < 0.01; and ***, P < 0.001.

We mapped *yq158* and the other 4 mutations to linkage group III (LGIII) and identified the affected gene, *Y71H2AM.9*, which encodes a protein that is homologous to the mammalian putative Zn^2+^ transporter SLC30A9 and the *E. coli* Zn^2+^ transporter Yiip (Fig. 1 E and Fig. S2) (Lu and Fu, 2007). We thus renamed the *Y71H2AM.9* gene *slc-30A9*. While *yq166*, *yq172* and *yq212* caused changes of amino acid residues, the *yq158* mutation caused a premature translation termination at Trp305 (W305stop) in the encoded protein. In addition, the *yq189* mutation affected a splicing site in *slc-30A9* mRNA (Fig. 1 E). Transgenic expression driven by the *slc-30A9* promoter of wild-type *slc-30A9*, but not mutant s*lc-30A9* containing the *yq158* mutation (W305stop), fully rescued the abnormal spherical mitochondria in *slc-30A9(yq158)* mutants to the wild-type level in multiple tissues (Fig. 1 F). Similar expression of GFP-fused SLC-30A9 (SLC-30A9::GFP), which is widely expressed in multiple tissues (Fig. S1 D), also rescued the enlargement of mitochondria labeled with TOMM-20::mCh (Figure 1G). SLC-30A9::GFP colocalized with TOMM-20::mCh (Fig. 1 G). When expressed in HeLa cells, GFP-fused *C. elegans* SLC-30A9 and GFP-fused human SLC30A9 were both found to localize to mitochondria stained with MitoTracker DR (Fig. S3, A and C). Taken together, these findings suggest that SLC-30A9 localizes to mitochondria and plays an essential role in maintaining mitochondrial structure and functions.

### SLC-30A9 is a mitochondrial Zn^2+^ exporter

By prediction, SLC-30A9 has a similar structure to that of the bacterial Yiip Zn^2+^ transporter (Lu and Fu, 2007) (Fig. 2 A). We thus examined whether SLC-30A9 has Zn^2+^ binding activity. In microscale thermophoresis (MST) assays, SLC-30A9-EGFP expressed in HEK293 cells bound with Zn^2+^, but not Ca^2+^, Mg^2+^ (Fig. 2 B, Fig. S3, A and B). SLC-30A9-EGFP probably bound with Cu^2+^ and Mn^2+^ with low affinity (Fig. S3, A and B). The Zn^2+^ binding activity localizes to the C-terminal part containing the cation-efflux domain, while the N-terminus has no obvious Zn^2+^ binding activity (Fig. 2 B). Mitochondrion-localized human SLC30A9-GFP similarly bound to Zn^2+^ (Fig. S3 C and D). We next mutated SLC-30A9 residues corresponding to those amino acids important for Zn^2+^ transporter activity in bacterial Yiip (Fig. 2 A and Fig. S2), and examined how this affected the rescuing effect on *slc-30A9(yq158)* mitochondria. When expressed under the control of the hypodermis-specific *col-19* promoter, wild-type SLC-30A9 fully rescued the mitochondrial defects in *slc-30A9(yq158)* hypodermis (Fig. 2 C). However, similarly expressed SLC-30A9 (H198A/D202A) or SLC-30A9(H402A) mutants had strongly reduced rescuing activity, and SLC-30A9(D221A) had no rescuing activity (Fig. 2 C). These results suggest that the Zn^2+^ transporter activity of SLC-30A9 is required for normal mitochondrial structure.

**Figure 2.**
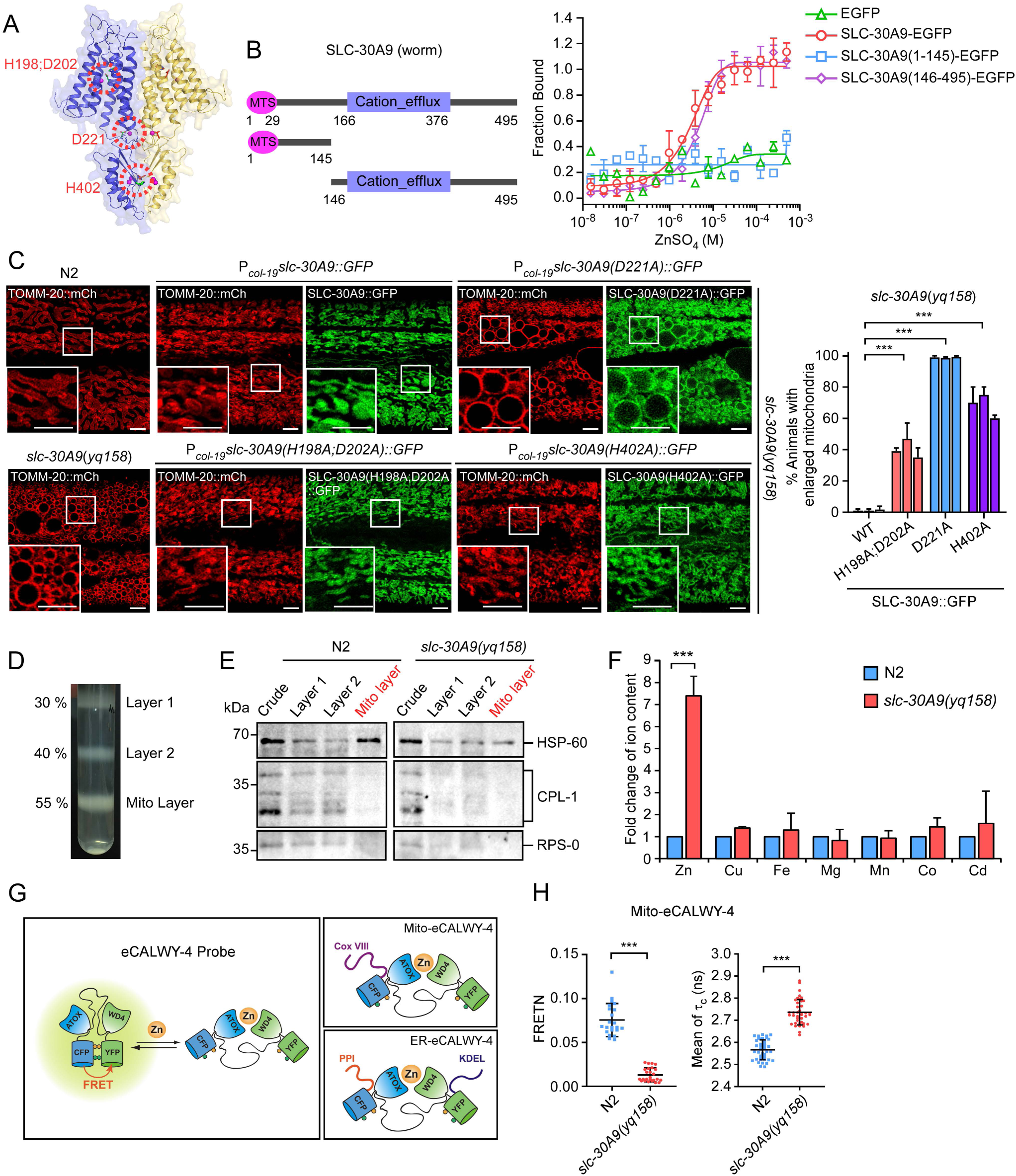
SLC-30A9 is a mitochondrial Zn^2+^ exporter. (**A**) Predicted structure of *C. elegans* SLC-30A9 bound to Zn^2+^ (magenta spheres). Ribbon representation of an SLC-30A9 homodimer (blue and yellow for each protomer). Residues important for Zn^2+^ binding are indicated in green and encircled in red. (**B**) Left: diagram of full-length and truncated SLC-30A9. Right: binding curve for Zn^2+^ with SLC-30A9-EGFP, SLC-30A9(1-145)-EGFP, SLC-30A9(146-495)-EGFP and EGFP expressed in HEK293 cells. (**C**) Images (left) and quantification (right) of the rescuing effects on *slc-30A9*(*yq158*) hypodermal mitochondria by GFP-fused SLC-30A9, SLC-30A9(H198A;D202A), SLC-30A9(D221A), and SLC-30A9(H402A) driven by the hypodermis-specific *col-19* promoter. Boxed regions are magnified (2.5×) in the insets. Bars, 5 μm. Three lines were analyzed for each transgene. ≥90 animals were scored for each transgenic line. Comparisons are between transgenes expressing wild-type (WT) and indicated SLC-30A9 mutant proteins. (**D**) Purification of mitochondria with sucrose density gradient centrifugation. (**E**) Western blotting of the samples in (**D**) with antibodies against mitochondrial HSP-60, lysosomal CPL-1, and ribosomal RPS-0. (**F**) Profiling of divalent cations in purified mitochondria from (**D**), measured with ICP-MS. (**G**) Schematic depiction of the eCALWY-4 Zn^2+^ sensors. Mitochondrial-targeted Mito-eCALWY-4 and ER-targeted ER-eCALWY-4 are expressed in the hypodermal cells of *C. elegans* to detect Zn^2+^ levels. (**H**) Normalized measure of FRET (FRETN) (left, measured by FRET) and the donor fluorescence lifetime (τ_c_) (right, measured by FLIM-FRET) of Mito-eCALWY-4 in the hypodermal cells of N2 and *slc-30A9*(*yq158*) animals. ≥30 animals were analyzed for each genotype. All experiments were performed in triplicate. For all quantifications, error bars represent SEM. *, P < 0.05; **, P < 0.01; and ***, P < 0.001.

We next examined mitochondrial Zn^2+^ levels in *slc-30A9* mutants. We purified mitochondria from both N2 and *slc-30A9(yq158)* mutants (Fig. 2, D and E) and assessed Zn^2+^ levels using ICP-MS (inductively coupled plasma mass spectrometry). In *slc-30A9(yq158)* mutants, the mitochondrial Zn^2+^ level was >7 times higher than in N2 animals (Fig. 2 F). Other divalent cations, including Cu^2+^, Fe^2+^, Mg^2+^, Mn^2+^, Co^2+^, and Cd^2+^, were not significantly changed (Fig. 2 F). Thus, loss of *slc-30A9* caused mitochondrial Zn^2+^ accumulation. To corroborate this result, we expressed a mitochondrion-localized Zn^2+^ sensor, Mito-eCALWY-4 (Chabosseau et al., 2014), to detect mitochondrial Zn^2+^ in *C. elegans* using FRET and FLIM-FRET assays (Gordon et al., 1998; Lin et al., 2015; Sun et al., 2011). In the absence of Zn^2+^, the eCALWY-4 sensor produces fluorescence resonance energy transfer, with a stronger normalized measure of FRET (FRETN) but a shorter florescence lifetime (τ_c_) of the donor fluorescent protein (CFP). When bound to Zn^2+^, eCALWY-4 generates a weaker FRETN while the fluorescence lifetime (τ_c_) of the donor is increased (Fig. 2 G) (Chabosseau et al., 2014; Gordon et al., 1998; Lin et al., 2015); (Sun et al., 2011). In the FRET and FLIM-FRET assays, *slc-30A9(yq158)* mutants had much weaker FRETN (measured by FRET) but significantly longer τ_c_ (measured by FLIM-FRET) of Mito-eCALWY-4 than N2 animals (Fig. 2 H). This suggests that Zn^2+^ accumulates in the mitochondria of *slc-30A9(yq158)* mutants. Collectively, the above results revealed that SLC-30A9 is a mitochondrial Zn^2+^ exporter required for mitochondrial Zn^2+^ efflux.

### Loss of *slc-30A9* impairs mitochondrial function, compromises animal development and shortens the life span

We next assessed how mitochondrial Zn^2+^ accumulation affects mitochondrial function and animal development. *slc-30A9* mutants had greatly reduced mitochondrial membrane potentials, and their ATP levels were strongly decreased (Fig. 3, A and B). Consistent with the reduction of ATP levels, the animal bending frequency of *slc-30A9* mutants was significantly lower than that of N2 animals (Fig. 3 C). Remarkably, *slc-30A9* mutants displayed strong mitochondrial unfolded protein responses (UPR^mt^), as evidenced by the strongly increased signals of GFP driven by the *hsp-6* promoter (P*_hsp-6_GFP*) in *slc-30A9* mutants (Lin et al., 2016; Nargund et al., 2015) (Fig. 3 D). In addition, *slc-30A9* mutants had far fewer progeny, indicating that loss of *slc-30A9* impairs fertility (Fig. 3 E). Around 30% of *slc-30A9* mutant embryos died (Fig. 3 F), and the development time from embryo to larval stage 4 (L4) of *slc-30A9* mutants was nearly 2 times that of N2 animals (Fig. 3 G). Moreover, the average life span of *slc-30A9* adults was decreased by >50% compared to N2 animals (Fig. 3 H). Taken together, these data suggest that, in the absence of *slc-30A9*, mitochondrial Zn^2+^ accumulation results in mitochondrial dysfunction and pleotropic defects of *C. elegans* development and physiology.

**Figure 3.**
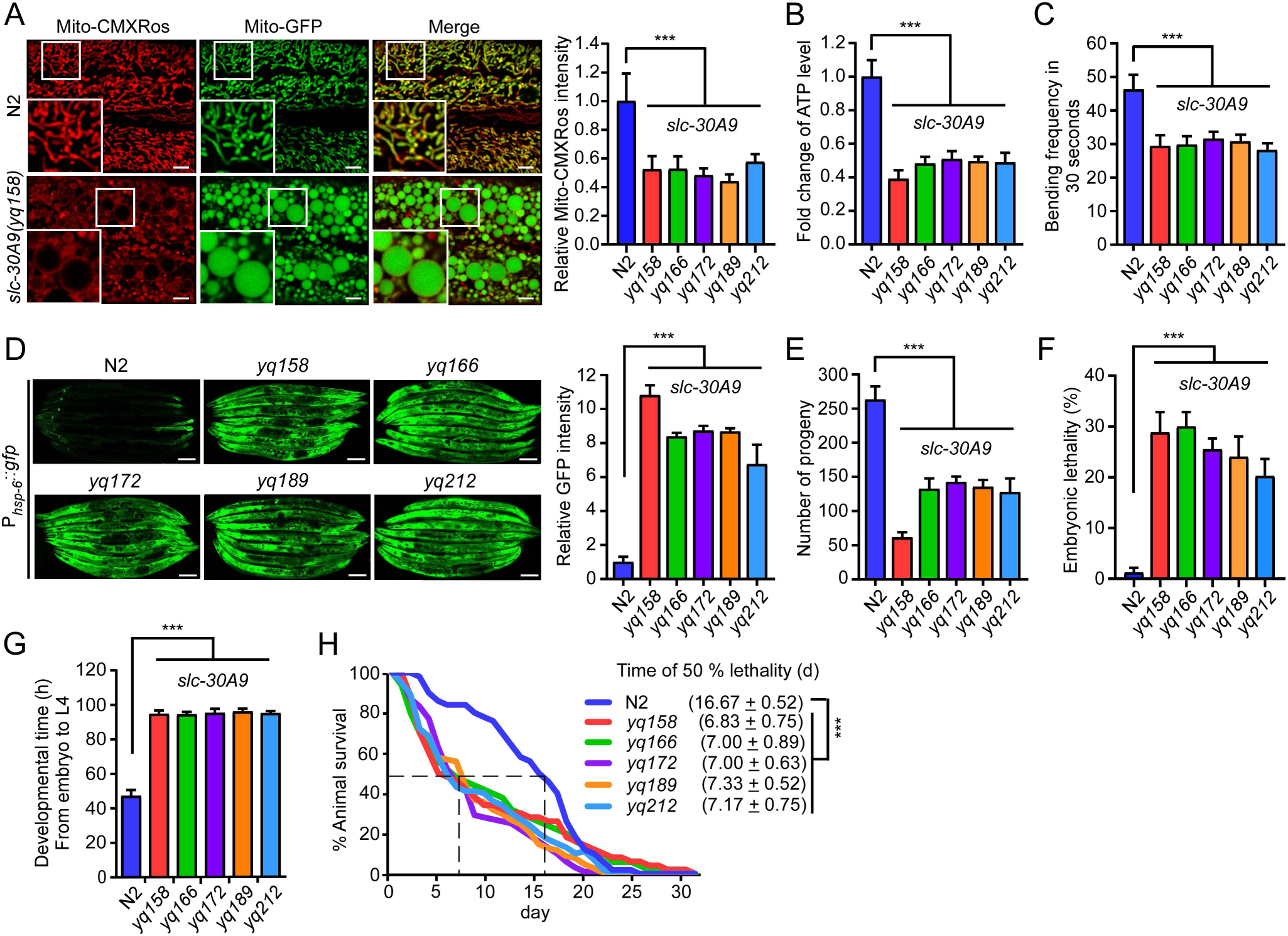
Loss of SLC-30A9 impairs mitochondrial functions, compromises animal development and shortens the life span. (**A**) Images (left) and quantification (right) of Mito-CMXRos staining in adult animals of N2 and *slc-30A9* mutants. Bars, 5 µm. 20 synchronized animals were analyzed for each genotype. (**B**) Fold change of ATP levels in adult animals of N2 and *slc-30A9* mutants. Data were from three independent experiments and normalized to the ATP levels in N2 animals. (**C**) Analysis of the bending frequency in 30 seconds of adult animals with the indicated genotype. 30 synchronized animals were analyzed for each genotype. (**D**) Representative images of UPR^mt^ indicated by the P*_hsp-6_GFP* reporter in adult N2 animals and *slc-30A9* mutants (left). Bars, 100 µm. Quantification of fluorescence intensities is shown on the right. Data were normalized to the GFP intensities in N2 animals. ≥90 animals were analyzed for each genotype. (**E** and **F**) Analysis of the progeny numbers (**E**) and embryonic lethality (**F**) in N2 and *slc-30A9* animals. 20 synchronized animals of each genotype were analyzed. (**G**) Analysis of the development time from embryo to L4 stage of N2 and *slc-30A9* mutant animals. 120 synchronized animals of each genotype were analyzed. (**H**) Survival curves of adult N2 animals and *slc-30A9* mutants. 120 animals were analyzed for each genotype. For all quantifications, error bars represent SEM. *, P < 0.05; **, P < 0.01; and ***, P < 0.001. Comparisons are between N2 and *slc-30A9* mutants.

### Mutations in *slc-25A25* suppress the abnormal mitochondrial enlargement in *slc-30A9* mutants

To investigate further how mitochondrial Zn^2+^ homeostasis is maintained, we carried out *slc-30A9* suppressor screens using two strategies. First, we mutagenized *slc-30A9(yq158)* animals and searched for mutants that displayed tubular mitochondria in adult hypodermis. Two mutants, *yq350* and *yq351*, were obtained. In double mutants of *yq350* or *yq351* with *slc-30A9(yq158)*, mitochondria exhibited predominantly tubular morphology in multiple tissues rather than the abnormally enlarged spherical mitochondria in *slc-30A9(yq158)* single mutants (Fig. 4 A). Second, we performed a suppressor screen on *slc-30A9(yq158)* expressing P*_hsp-6_GFP* to look for mutants with greatly reduced UPR^mt^ signals. Three mutants, *yq371*, *yq372*, and *yq373*, had strongly reduced GFP (UPR^mt^) signals when doubled with *slc-30A9(yq158)* (Fig. 4 B). Using SNP mapping and sequencing, we unexpectedly found that all 5 of these mutants affected the same gene, *F17E5.2*, on linkage group X (LGX). Within the *F17E5.2* gene, *yq371* caused a mis-splicing mutation, while the other 4 mutations changed amino acid residues in the encoded protein (Fig. 4 C). In addition, we used the CRISPR/Cas9 system to generate another mutant, *yq406,* which carried an early stop-codon mutation in *F17E5.2* (Fig. 4 C). Double mutants of *yq406* with *slc-30A9(yq158)* had tubular mitochondria and strongly reduced UPR^mt^, confirming that loss of *F17E5.2* suppressed the abnormal enlargement of mitochondria in the absence of *slc-30A9* (Fig. 4, A and B).

**Figure 4.**
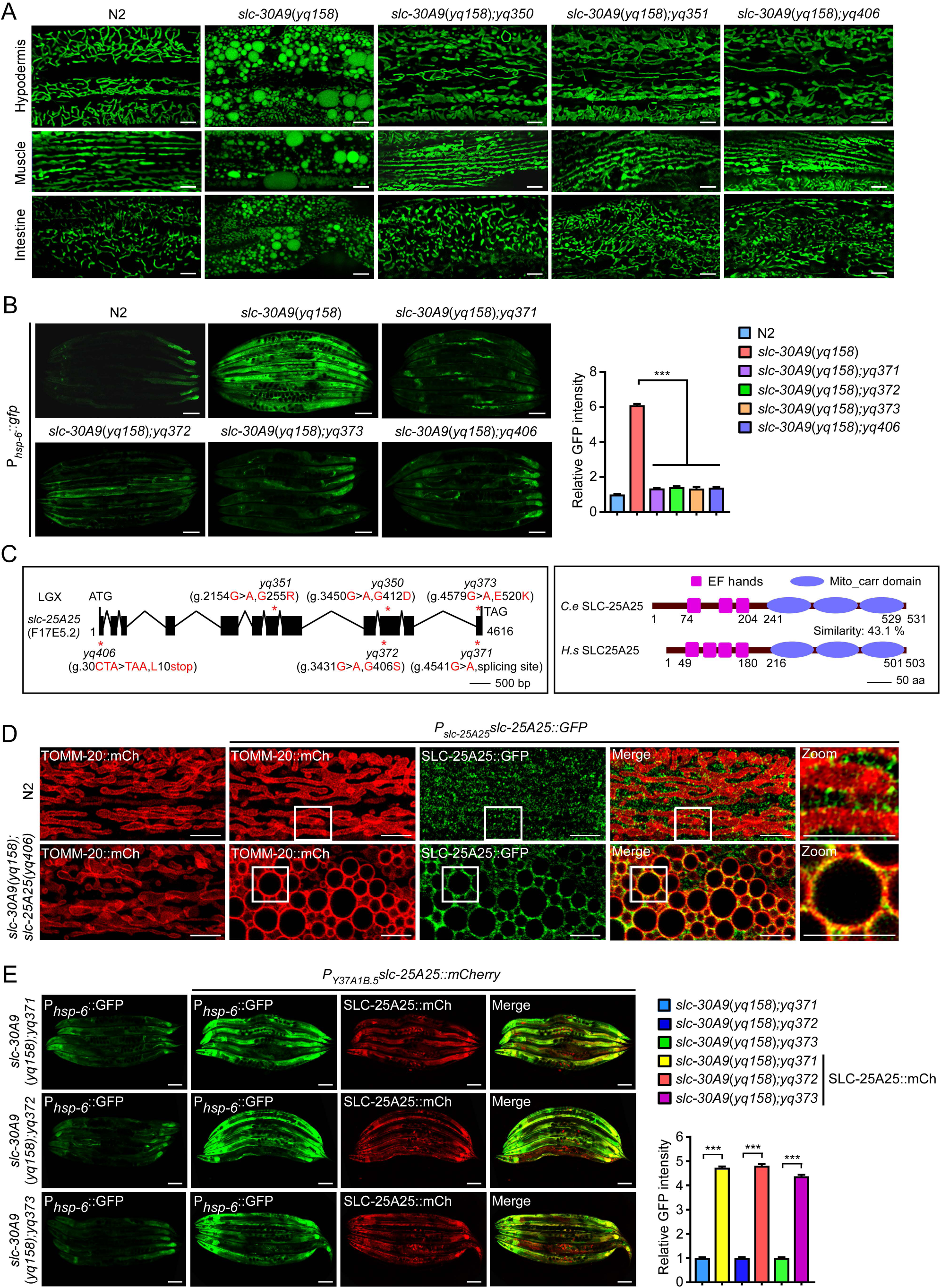
Loss *slc-25A25* suppresses mitochondrial defects in *slc-30A9* mutants. (**A**) Representative images of mitochondria in hypodermal, muscle and intestinal cells in N2 animals, *slc-30A9*(*yq158*) single mutants, and double mutants of *slc-30A9*(*yq158*) with 2 identified suppressors (*yq350* and *yq351*) and the CRISPR/Cas9-engineered mutation *yq406*. Bars, 5 μm. (**B**) Representative images (left) and quantification (right) of UPR^mt^ in N2 animals, *slc-30A9*(*yq158*) single mutants, and double mutants of *slc-30A9*(*yq158*) with 3 identified suppressors (*yq371, yq372* and *yq373*) and the CRISPR/Cas9-engineered mutation *yq406*. Bars, 100 μm. ≥90 animals were analyzed for each genotype. Data were from three independent experiments and normalized to the GFP intensities in N2 animals. (**C**) Schematic representation of the *slc-25A25* gene (left) and the encoded protein (right). Filled boxes represent exons and thin lines indicate introns. The point mutations of *slc-25A25* are indicated with asterisks. The EF hands and Mito-carrier domain are indicated in magenta and blue, respectively. Amino acid similarity is shown between *C. elegans* SLC-25A25 and human SLC25A25. (**D**) Images of the rescuing effects on *slc-30A9*(*yq158*);*slc-25A25*(*yq406*) hypodermal mitochondria of GFP-fused SLC-25A25 driven by the *slc-25A25* promoter. Bars, 5 µm. Boxed regions are magnified (2.5×) in the insets. Bars, 5 µm. (**E**) Images (left) and quantification (right) of the rescuing effects of the *Y37A1B.5* promoter-driven SLC-25A25::mCh on the UPR^mt^ in double mutants of *slc-30A9*(*yq158*) with *slc-25A25*(*yq371*), *slc-25A25*(*yq372*), and *slc-25A25*(*yq373*). Bars, 100 µm. ≥90 animals analyzed for each genotype. For all quantifications, *, P < 0.05; **, P < 0.01; and ***, P < 0.001. Error bars represent SEM.

*F17E5.2* encodes a homolog of mammalian SLC25A25/SCaMC-2 (short calcium-dependent mitochondrial carrier 2), a Ca^2+^-activated mitochondrial Mg^2+^-ATP transporter with 4 EF-hands (EFhs) and 3 mitochondrial carrier domains (Fig. 4 C and Fig. S4 A) (del Arco and Satrustegui, 2004; Fiermonte et al., 2004; Joyal and Aprille, 1992; Satrustegui et al., 2007; Yang et al., 2014). By predication, F17E5.2 only has 3 EFhs (Fig. 4 C and Fig. S4 A). *F17E5.2* was thus renamed *slc-25A25*. Transgenic expression of *slc-25A25* under the control of its own promoter or the hypodermal *Y37A1B.5* promoter restored the abnormal mitochondrial enlargement as well as the strong UPR^mt^ in *slc-30A9(yq158);slc-25A25(yq406)* double mutants (Fig. 4, D and E). These findings provide further evidence that *slc-25A25* is responsible for rescue of *slc-30A9*-induced mitochondrial defects. Like SLC-30A9, SLC-25A25 expression was seen in multiple tissues (Fig. S4 B). In addition, mitochondrion-localized SLC-25A25::GFP was observed in N2 animals and was more evident in *slc-30A9;slc-25A25* double mutants (Fig. 4 D), which suggests that SLC-25A25 likely functions on mitochondria.

### SLC-25A25 is important for normal mitochondrial structure and function

We next investigated whether *slc-25A25* mutations caused mitochondrial defects. Although loss of *slc-25A25* suppressed the abnormal enlargement of spherical mitochondria in *slc-30A9* animals, the mitochondria in *slc-25A25(yq406)* single mutants appeared as abnormally flattened tubules in multiple tissues compared with N2 animals (Fig. 5 A). Using TEM, we found that mitochondria in *slc-25A25(yq406)* hypodermal cells were morphologically abnormal compared with those in N2 animals (Fig. 5 B). In *slc-25A25(yq406)* muscle cells, mitochondria contained vacuolar structures (Fig. 5 B). *slc-25A25* mutants did not have an obvious UPR^mt^ (Fig. 5 C), consistent with the finding that loss of *slc-25A25* suppressed the UPR^mt^ induced by *slc-30A9* loss of function (Fig. 4 B).

**Figure 5.**
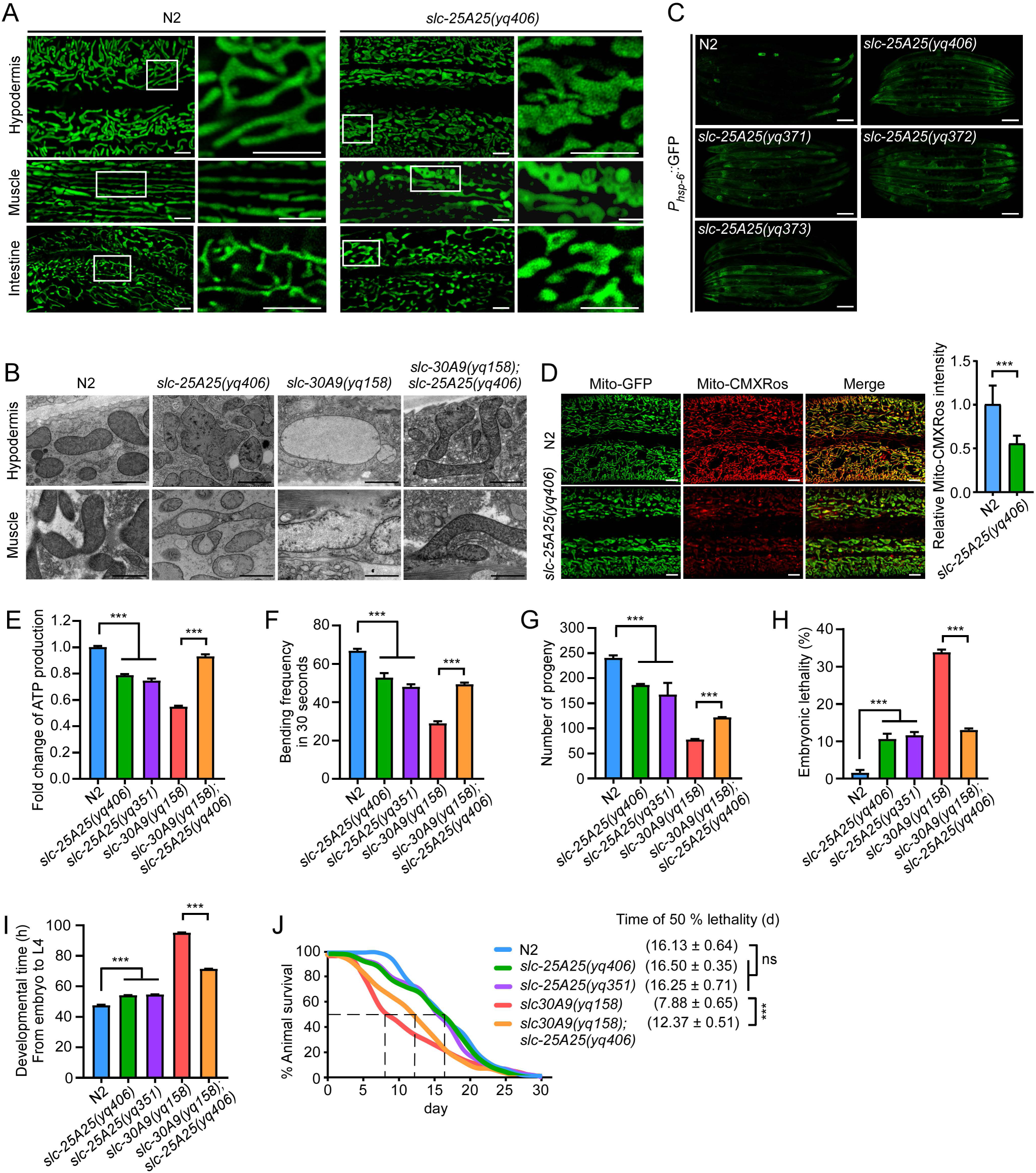
SLC-25A25 is important for normal mitochondrial structure and functions. (**A**) Representative images of mitochondria in hypodermal, muscle and intestinal cells in N2 and *slc-25A25*(*yq406*) animals. For each genotype, boxed regions are magnified (4 ×) and shown at the right. Bars, 5 μm. (**B**) Representative TEM images of mitochondria in hypodermal and muscle cells in N2, *slc-25A25*(*yq406*), *slc-30A9*(*yq158*), and *slc-30A9*(*yq158*);*slc-25A25*(*yq406*) animals. Bars, 1 μm. (**C**) Representative images of the UPR^mt^ (P*_hsp-6_GFP*) in N2 animals and the indicated *slc-25A25* mutants. Bars, 100 μm. (**D**) Images (left) and quantification (right) of Mito-CMXRos intensity in adult N2 and *slc-25A25*(*yq406*) animals. 20 synchronized animals were analyzed for each genotype. (**E**) Fold change of ATP levels in N2, *slc-25A25*(*yq406*), *slc-25A25*(*yq351*), *slc-30A9*(*yq158*), and *slc-30A9*(*yq158*);*slc-25A25*(*yq406*) adults. Data were from three independent experiments and normalized to the ATP levels in N2 animals. (**F**) Bending frequencies of N2, *slc-25A25*(*yq406*), *slc-25A25*(*yq351*), *slc-30A9*(*yq158*), and *slc-30A9*(*yq158*);*slc-25A25*(*yq406*) adults. 30 synchronized animals were analyzed for each genotype. (**G** and **H**) Analysis of the progeny numbers (**G**) and embryonic lethality (**H**) in adult N2, *slc-25A25*(*yq406*), *slc-25A25*(*yq351*), *slc-30A9*(*yq158*), and *slc-30A9*(*yq158*);*slc-25A25*(*yq406*) animals. 20 synchronized animals of each genotype were analyzed. (**I**) Analysis of the development time from embryo to L4 stage of N2, *slc-25A25*(*yq406*), *slc-25A25*(*yq351*), *slc-30A9*(*yq158*), and *slc-30A9*(*yq158*);*slc-25A25*(*yq406*) animals. 120 synchronized animals of each genotype were analyzed. (**J**) Survival curves of adult N2, *slc-25A25*(*yq406*), *slc-25A25*(*yq351*), *slc-30A9*(*yq158*), and *slc-30A9*(*yq158*);*slc-25A25*(*yq406*) animals. 120 animals were analyzed for each genotype. For all quantifications, *, P < 0.05; **, P < 0.01; and ***, P < 0.001. Error bars represent SEM.

We further analyzed the effect of *slc-25A25* mutations on mitochondrial functions and animal development. The *slc-25A25* mutants *yq406* and *yq351* exhibited significantly reduced mitochondrial membrane potential and ATP levels, and the animal bending rates were significantly decreased (Fig, 5, D-F). In addition, *slc-25A25* mutants generated fewer progeny, displayed about 10% embryonic lethality, and had slower development (Fig. 5, G-I). Nevertheless, *slc-25A25* adults appeared to have a similar life span to N2 animals (Fig. 5 J). Importantly, compared with *slc-30A9(yq158)* single mutants, double mutants of *slc-25A25 (yq406)* with *slc-30A9(yq158)* exhibited nearly normal mitochondrial structures (Fig. 5 B), enhanced ATP production and motor ability (Fig. 5, E and F), more progeny, decreased embryonic lethality, and faster development (Fig. 5, G-I). The median life span of the double mutants was significantly enhanced compared with *slc-30A9(yq158)* single mutants (Fig. 5 J). Taken together, these results suggest that SLC-25A25 is important for normal mitochondrial structure and functions, and that *slc-25A25* functions to antagonize *slc-30A9*.

### SLC-25A25 binds with Zn^2+^ and is important for mitochondrial Zn^2+^ accumulation in *slc-30A9* mutants

Using the Mito-eCALWY-4 Zn^2+^ sensor and FLIM-FRET, we assessed whether loss of *slc-25A25* suppressed the mitochondrial Zn^2+^ accumulation caused by loss of *slc-30A9*. Compared with N2 animals, the donor fluorescence lifetime (τ_c_) of Mito-eCALWY-4 in *slc-30A9(yq158)* animals was strongly increased (Figs. 2 H and 6 A), indicating the elevated mitochondrial Zn^2+^ level. In *slc-30A9(yq158);slc-25A25(yq406)* double mutants, however, the donor fluorescence lifetime (τ_c_) in Mito-eCALWY-4 was significantly reduced (Fig. 6 A). Thus, loss of *slc-25A25* inhibited the mitochondrial Zn^2+^ accumulation in *slc-30A9* mutants, which suggests that SLC-25A25 regulates mitochondrial Zn^2+^ levels. To investigate this further, we examined whether SLC-25A25 binds with Zn^2+^. In MST assays, purified recombinant worm SLC-25A25-EGFP bound with Zn^2+^ (*K*d 2.40±0.27 μM) (Fig. 6 B). Likewise, recombinant human SLC25A25-EGFP bound with Zn^2+^ (*K*d 1.87±0.31 μM). EGFP-tagged *C. elegans* SLC-25A25 and human SLC25A25 both bound Ca^2+^ with similar affinity (Fig. 6, B and C). As a negative control, recombinant EGFP had no obvious Zn^2+^- or Ca^2+^- binding activity (Fig. 6 D). These findings suggest that both *C. elegans* SLC-25A25 and human SLC25A25 can bind with Zn^2+^ in addition to Ca^2+^. Notably, transgenic expression of GFP-fused human SLC25A25 driven by the *C. elegans slc-25A25* promoter restored the mitochondrial enlargement in *slc-30A9(yq158);slc-25A25(yq406)* double mutants (Fig. 6 E), suggesting an evolutionarily conserved function of SLC25A25.Interestingly, double mutants of *slc-30A9(yq158)* with *mcu-1(ju1154)*, a strong loss-of-function mutant of the worm mitochondrial calcium uniporter (MCU), exhibited a similar mitochondrial abnormality to *slc-30A9(yq158)* single mutants (Fig. 6 F). This suggests that MCU is not responsible for the mitochondrial enlargement induced by loss of *slc-30A9*.

**Figure 6.**
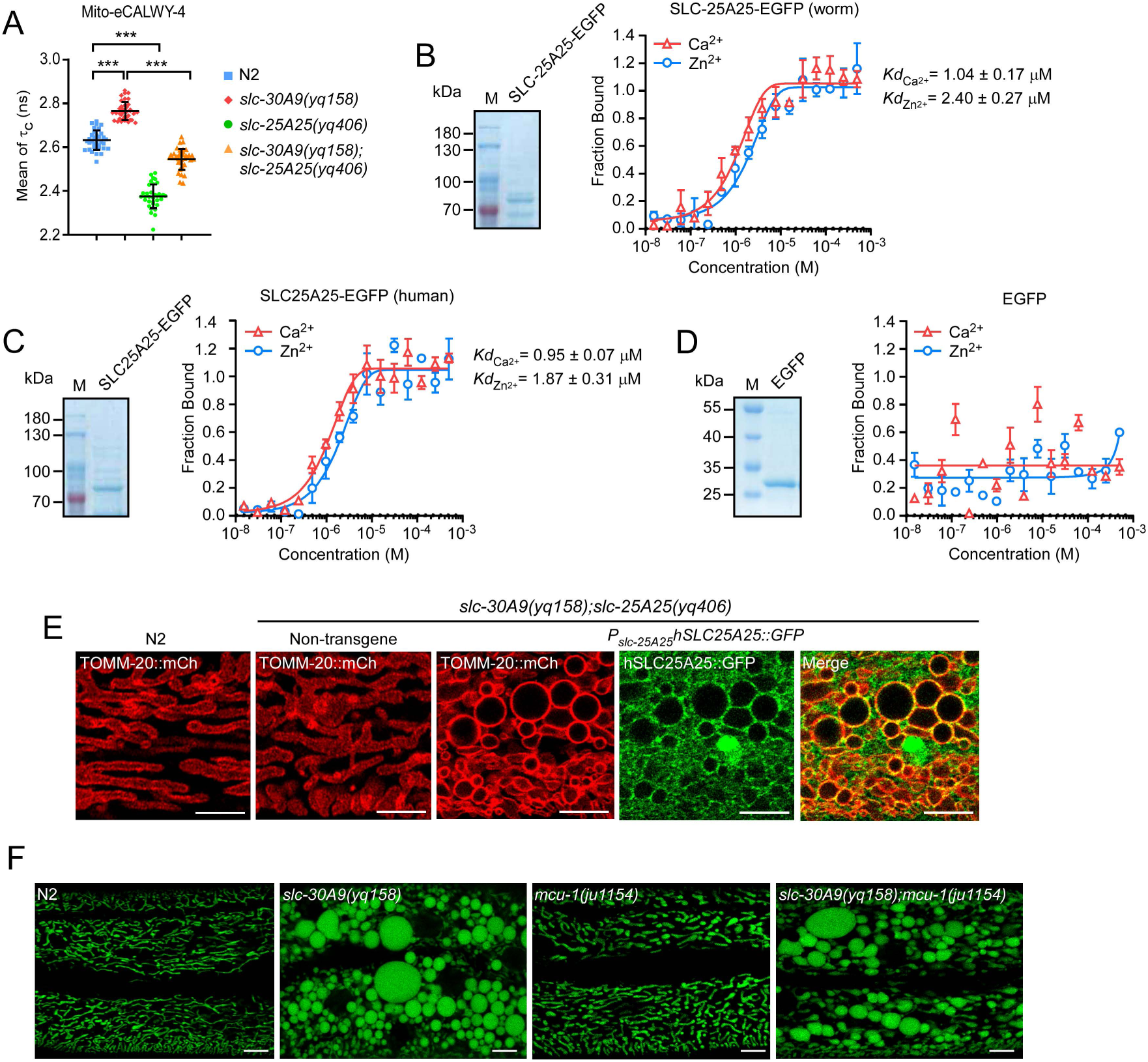
SLC-25A25 binds with Zn^2+^ and is required for mitochondrial Zn^2+^ accumulation in *slc-30A9* mutants. (**A**) The donor fluorescence lifetime (τ_c_) of Mito-eCALWY-4 in the hypodermal cells of N2, *slc-30A9*(*yq158*), *slc-25A25*(*yq406*), and *slc-30A9*(*yq158*);*slc-25A25*(*yq406*) animals. ≥30 animals were analyzed for each genotype. Data were derived from three independent experiments. (**B**-**D**) Binding curves of Zn^2+^ and Ca^2+^ with recombinant *C. elegans* SLC-25A25-EGFP (**B**), human SLC25A25-EGFP (**C**), and EGFP (**D**). Purified recombinant proteins are shown on the left in each panel. (**E**) Images representing the rescuing effects of human SLC25A25-GFP driven by the *C. elegans slc-25A25* promoter on *slc-30A9*(*yq158*);*slc-25A25*(*yq406*) hypodermal mitochondria labeled with TOMM-20::mCh. Bars, 5 µm. (**F**) Representative images of hypodermal mitochondria in N2, *mcu-1(ju1154)*, *slc-30A9(yq158)*, and *slc-30A9(yq158);mcu-1(ju1154)* animals. Bars, 5 µm. For all quantifications, *, P < 0.05; **, P < 0.01; and ***, P < 0.001. NS, not significant. Error bars represent SEM.

### SLC-25A25/SLC25A25 overexpression leads to mitochondrial enlargement and Zn^2+^ elevation

To investigate whether *C. elegans* SLC-25A25 and human SLC25A25 promote mitochondrial Zn^2+^ import, we generated HeLa cell lines expressing EGFP-tagged SLC-25A25 (worm) or SLC25A25 (human). We then purified mitochondria and measured divalent ions using ICP-MS. Our results indicated that mitochondrial Zn^2+^ levels were strongly increased in cells expressing EGFP-SLC-25A25 (worm) or EGFP-SLC25A25 (human) (Fig. 7, A and B). No obvious changes were detected for other divalent ions including Cu^2+^, Fe^2+^, Mg^2+^, Mn^2+^, Co^2+^ and Ca^2+^ (Fig. 7, A and B). Compared with the control cells, the cells expressing worm SLC-25A25 or human SLC25A25 had significantly enlarged mitochondria, which were further enlarged by supplementation of the culture medium with 0.1 mM Zn^2+^ (Fig. 7 C). Furthermore, the enlarged mitochondria were strongly positive for the Zn^2+^ indicator Zinpyr-1, which indicates that they contained elevated levels of Zn^2+^ compared with the mitochondria in control cells (Fig. 7 D). Collectively, these results provided further evidence that SLC-25A25 promotes mitochondrial import of Zn^2+^.

**Figure 7.**
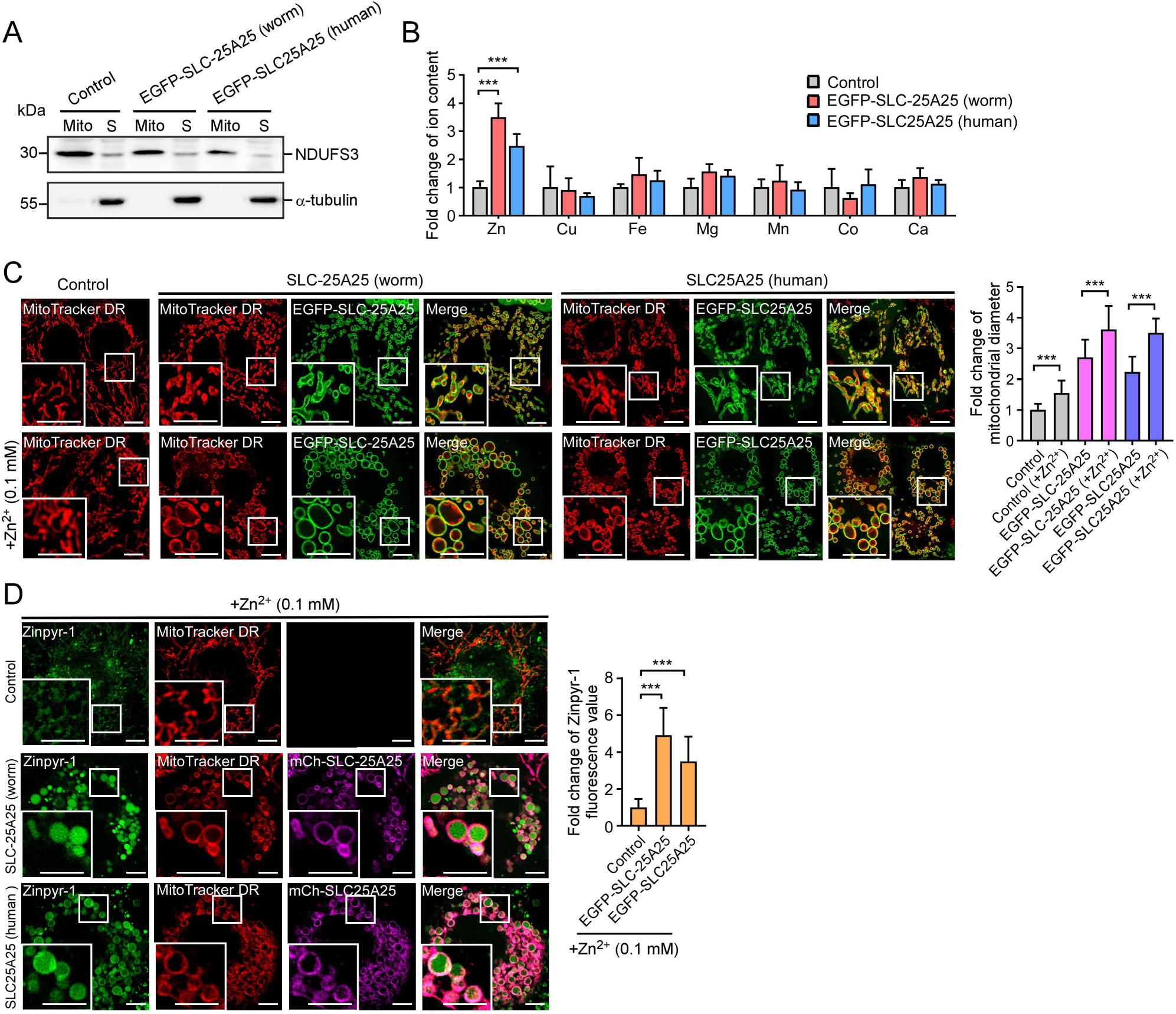
Ectopic expression of SLC-25A25 causes abnormal mitochondrial enlargement in HeLa cells. (**A**) Western blotting analysis of purified mitochondria using an antibody against mitochondrial NDUFS3. Mitochondria were purified from cell lines stably expressing EGFP-SLC-25A25 (worm) or EGFP-SLC25A25 (human). Cytoplasmic α-tubulin was used as control. Mito: mitochondria. S: supernatant. (**B**) Profiling of divalent cations in mitochondria purified from cell lines stably expressing EGFP-SLC-25A25 (worm) or EGFP-SLC25A25 (human), measured with ICP-MS. Levels of individual divalent cations were normalized to those in the control cells. (**C**) Representative images of mitochondria (left) and quantification (right) of mitochondrial diameters in HeLa cells expressing EGFP-SLC-25A25 (worm) or EGFP-SLC25A25 (human). Cells were cultured without or with additional Zn^2+^ (0.1 mM). Mitochondria are labeled with MitoTracker DR. Boxed regions are magnified (2 ×) in the bottom left. Bars, 2 µm. ≥ 40 mitochondria from ≥ 7 cells were analyzed for each treatment. (**D**) HeLa cells expressing EGFP-SLC-25A25 (worm) or EGFP-SLC25A25 (human) were cultured without or with additional Zn^2+^ (0.1 mM) and treated with the fluorogenic Zn^2+^ reporter Zinpyr-1. Mitochondria are labeled with MitoTracker DR. In the images (left), boxed regions are magnified (2×) at the bottom left. Bars, 2 µm. Quantifications of Zinpyr-1 fluorescence are shown on the right. ≥ 40 mitochondria from ≥ 7 cells were analyzed for each treatment. For all quantifications, *, P < 0.05; **, P < 0.01; and ***, P < 0.001. Error bars represent SEM.

### Inhibiting the ER-localized Zn^2+^ transporter SLC-30A5 suppresses the mitochondrial Zn^2+^ accumulation in *slc-30A9* mutants

Because cellular Zn^2+^ is compartmentalized (Colvin et al., 2010), we sought to determine the location of the Zn^2+^ pool that provides the Zn^2+^ for mitochondria. To do this, we performed a candidate RNAi screen to knock down putative *C. elegans* Zn^2+^ transporters (Table S2). RNAi of *slc-30A5*/*ZnT5* strongly inhibited the abnormal enlargement of spherical mitochondria in *slc-30A9(yq158)* mutants (Fig. 8 A). TEM analysis revealed that *slc-30A5* RNAi-treated *slc-30A9(yq158)* animals exhibited tubular mitochondria in hypodermal cells, in contrast to the abnormally enlarged mitochondria in control RNAi-treated *slc-30A9(yq158)* mutants (Fig. 8 B). SLC-30A5/Znt-5 is predicted to be a Zn^2+^ transporter localized to the endoplasmic reticulum (ER). In line with this, EGFP-fused SLC-30A5 (EGFP-SLC-30A5) ectopically expressed in HeLa cells co-localized with the ER marker KDEL-mCh, and bound with Zn^2+^ in MST assays (Fig. 8 C). In *C. elegans*, transgenic SLC-30A5::GFP under the control of the *slc-30A5* promoter was found to co-localize with the ER marker SPCS-1 tagged with mCherry (SPCS-1::mCh) (Fig. 8 D). In FLIM-FRET assays, *slc-30A5* RNAi treatment significantly reduced Zn^2+^ levels in both ER and mitochondria in N2 and *slc-30A9(yq158)* animals, as evidenced by the reduction in the fluorescence lifetime of CFP (τ_c_) of ER- and Mito-eCALWY-4 probes (Fig. 8 E). These findings suggest that mitochondrial Zn^2+^ is imported from the ER. Using the ER-eCALWY-4 Zn^2+^ sensor, we found that *slc-25A25(yq406)* animals as well as *slc-30A9(yq158);slc-25A25(yq406)* double mutants had significantly higher ER Zn^2+^ levels (higher τ_c_ from the ER-eCALWY probe) than the wild type (Fig. 8 F). Furthermore, in both *slc-25A25(yq406)* single mutants and *slc-30A9(yq158);slc-25A25(yq406)* double mutants, *slc-30A5* RNAi significantly reduced the Zn^2+^ levels in both the ER and mitochondria, as evidenced by the decreased τ_c_ from the ER- and Mito-eCALWY-4 probes (Fig. 8 G). Thus, decreasing SLC-30A5-mediated Zn^2+^ uptake into the ER reduced the mitochondrial Zn^2+^ levels, suggesting that the ER likely serves as the Zn^2+^ pool for mitochondrial Zn^2+^ import.

**Figure 8.**
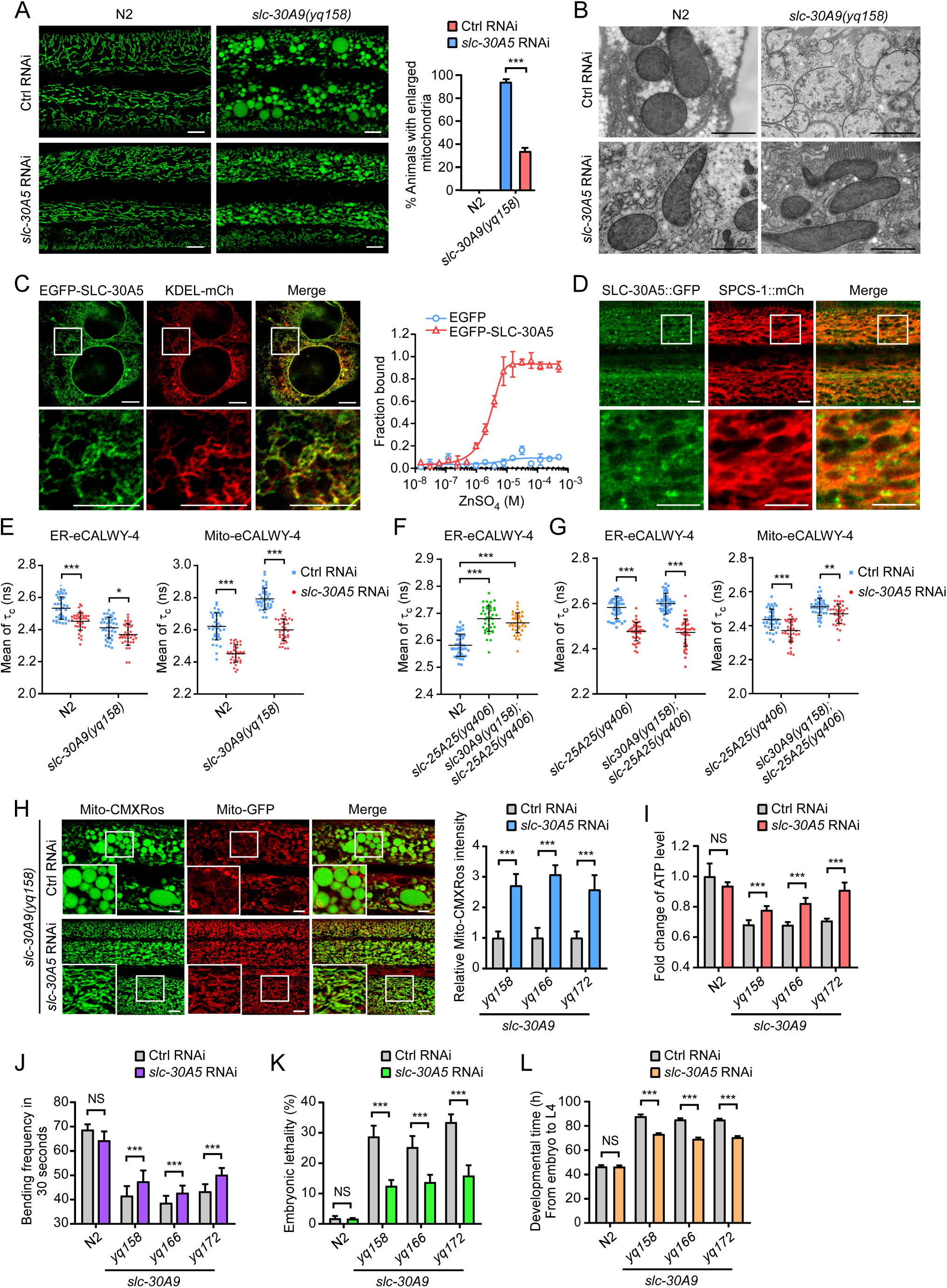
Inactivation of *slc-30A5* suppresses the mitochondrial abnormality in *slc-30A9* mutants. (**A**) Images (left) and quantification (right) of hypodermal mitochondria in control (Ctrl) RNAi- and *slc-30A5* RNAi-treated N2 and *slc-30A9*(*yq158*) animals. Bars, 5 µm. ≥90 animals were analyzed for each genotype. (**B**) Representative TEM images of mitochondria in hypodermal cells of Ctrl RNAi- and *slc-30A5* RNAi-treated N2 and *slc-30A9*(*yq158*) animals. Bars, 1 µm. (**C**) Left: localization of *C. elegans* EGFP*-*SLC-30A5 ectopically expressed in HeLa cells. Boxed regions in the top images are magnified (3.5×) and shown underneath. Bars, 1 µm. Right: binding curve for Zn^2+^ with *C. elegans* EGFP-SLC-30A5 expressed in HEK293 cells, measured with MST assays. EGFP was used as the negative control. (**D**) Localization of *C. elegans* SLC-30A5::GFP (P*_slc-30A5_slc-30A5::GFP*) in adult N2 animals. Boxed regions in the top images are magnified (3×) and shown underneath. Bars, 1 µm. (**E**) Donor fluorescence lifetime (τ_c_) of ER-eCALWY-4 (left) and Mito-eCALWY-4 (right) in the hypodermis of Ctrl RNAi- and *slc-30A5* RNAi-treated N2 and *slc-30A9(yq158)* animals. 30 synchronized animals were analyzed for each genotype. Data (mean ± SEM) were derived from three independent experiments. (**F**) Donor fluorescence lifetime (τ_c_) of ER-eCALWY-4 in the hypodermis of N2, *slc-25A25(yq406)*, and *slc-30A9(yq158);slc-25A25(yq406)* animals. 30 synchronized animals were analyzed for each genotype. Data (mean ± SEM) were derived from three independent experiments. (**G**) Donor fluorescence lifetime (τ_c_) of ER-eCALWY-4 (left) and Mito-eCALWY-4 (right) in the hypodermis of Ctrl RNAi- and *slc-30A5* RNAi-treated *slc-25A25(yq406)* and *slc-30A9(yq158);slc-25A25(yq406)* animals. 30 synchronized animals were analyzed for each genotype. Data (mean ± SEM) were derived from three independent experiments. (**H**) Left: Images of Mito-CMXRos staining in Ctrl RNAi- and *slc-30A5* RNAi-treated *slc-30A9*(*yq158*) animals. Boxed regions are magnified (2×) at the bottom left. Bars, 5 µm. Right: Quantification of Mito-CMXRos staining in the indicated *slc-30A9* mutants treated with Ctrl or *slc-30A5* RNAi. 20 synchronized animals (24 h post L4) were analyzed for each genotype. (**I**) Fold change of ATP levels in adult N2 animals and *slc-30A9* mutants treated with Ctrl RNAi or *slc-30A5* RNAi. Data were from three independent experiments and normalized to the ATP levels in N2 animals. (**J**) The bending frequency of Ctrl RNAi- and *slc-30A5* RNAi-treated animals with the indicated genotype. 30 synchronized animals were analyzed for each genotype. (**K**) The embryonic lethality of N2 and the indicated *slc-30A9* mutants treated with Ctrl RNAi or *slc-30A5* RNAi. 20 animals of each genotype were analyzed. (**L**) The development time from the embryo to L4-stage of N2 and *slc-30A9* mutants treated with Ctrl RNAi or *slc-30A5* RNAi. 120 synchronized animals of each genotype were analyzed.

We investigated the physiological requirement for the ER Zn^2+^ pool in mitochondrial functions. RNAi of *slc-30A5* restored the mitochondrial structure in *slc-39A9(yq158)* mutants (Fig. 8 B). Furthermore, mitochondrial membrane potentials were greatly enhanced by *slc-30A5* RNAi in *yq158* and other mutants of *slc-30A9*, as indicated by Mito-CMXRos staining (Fig. 8 H). In *slc-30A9* mutant animals, the ATP levels and the body bending frequencies were also significantly improved by *slc-30A5* RNAi (Fig. 8 I and J). In addition, *slc-30A5* RNAi significantly reduced the embryonic lethality and shortened the larval developmental time of *slc-30A9* mutants (Fig. 8 K and L). These results suggest that reducing SLC-30A5-mediated ER Zn^2+^ import ameliorated the structural and functional mitochondrial defects in *slc-30A9* mutants.

### *ZnT5* inactivation ameliorates SLC-25A25-related mitochondrial defects

To consolidate the findings that the ER provides the Zn^2+^ pool for mitochondrial Zn^2+^, we treated HeLa cells expressing worm SLC-25A25 or human SLC25A25 with siRNA (small RNA interference) of ZnT5, the human ortholog of *C. elegans* SLC-30A5. While *ZnT5* siRNA did not obviously change mitochondrial morphology in control cells, the Zn^2+^-induced mitochondrial enlargement in cells expressing GFP-SLC-25A25 (worm) or GFP-SLC25A25 (human) was strongly inhibited (Fig. 9, A and B). Moreover, *ZnT5* siRNA significantly reduced the elevation in mitochondrial Zn^2+^ levels, as evidenced by the reduced mitochondrial fluorescence intensity of the Zn^2+^ reporter Zinpyr-1 (Fig. 9 C). These results indicated that SLC-25A25/SLC25A25-mediated mitochondrial Zn^2+^ import depends on ZnT5-mediated ER Zn^2+^ uptake, thus providing further evidence that the ER serves as the Zn^2+^ pool for mitochondrial Zn^2+^ import. Supporting this conclusion, SLC-25A25::GFP (worm) and GFP-SLC25A25 (human) were found at mitochondrion-ER contacts in worm and HeLa cells, respectively (Fig. 9, D and E). In *slc-30A9(yq158);slc-25A25(yq406)* double mutants, the functional SLC-25A25::GFP was even more obviously enriched at mitochondrion-ER contacts (Fig. 9 D). Taken together, these findings suggest that the ER contains the Zn^2+^ pool for mitochondrial Zn^2+^ import, and that mitochondrial import of Zn^2+^ from the ER is probably achieved through mitochondrion-ER contacts.

**Figure 9.**
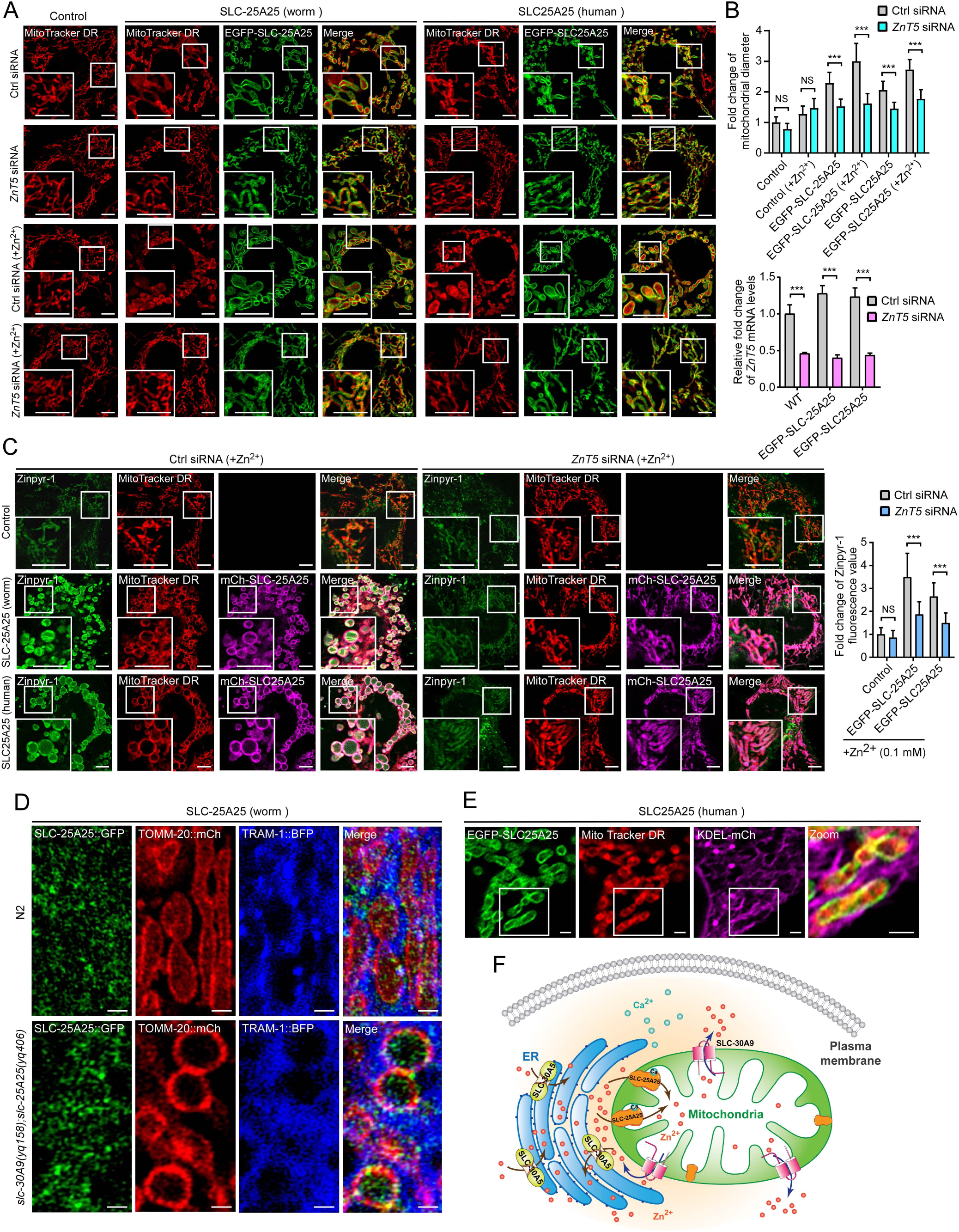
siRNA knockdown of human *SLC30A5/ZnT-5* ameliorates mitochondrial defects in cells ectopically expressing *C. elegans SLC-30A5* or human SLC25A25. **(A)** Representative images and quantification (right) of mitochondria in cells expressing EGFP-SLC-25A25 (worm) or EGFP-SLC25A25 (human) treated with Ctrl siRNA or *ZnT5* siRNA with or without additional Zn^2+^ (0.1 mM). Mitochondria were labeled with MitoTracker DR. Boxed regions are magnified (2×) in the bottom left. Bars, 2 µm. (**B**) Top: Quantification of mitochondrial diameters in cells as shown in **(A)**. ≥ 40 mitochondria from ≥ 7 cells were analyzed for each treatment. Bottom: mRNA levels of *ZnT5* in Ctrl siRNA- and *ZnT5* siRNA-treated control HeLa cells and cells expressing EGFP-SLC-25A25 (worm) or EGFP-SLC25A25 (human). (**C**) Images (left and middle) and quantification (right) of mitochondrial Zn^2+^ in HeLa cells expressing EGFP-SLC-25A25 (worm) or EGFP-SLC25A25 (human) treated with Ctrl siRNA or *ZnT5* siRNA with additional Zn^2+^ (0.1 mM). Mitochondrial Zn^2+^ was detected with the fluorogenic reporter Zinpyr-1. Mitochondria were labeled with MitoTracker DR. Boxed regions are magnified (2×) in bottom left. Bars, 2 µm. ≥ 40 mitochondria from ≥ 7 cells were analyzed for each treatment. (**D**) Localization of *C. elegans* SLC-25A25::GFP with TOMM-20::mCh and the ER marker TRAM-1::BFP in hypodermal cells of N2 and *slc-30A9*(*yq158*)*;slc-25A25(yq406)* animals. Bars, 1 µm. (**E**) Localization of human EGFP-SLC25A25 with MitoTracker DR and KDEL-mCh in HeLa cells. Boxed regions are zoomed (3×) and merged in the right-most image. Bars, 2 µm. (**F**) Schematic summary of the mechanisms that control mitochondrial Zn^2+^ transport. For all quantifications, *, P < 0.05; **, P < 0.01; and ***, P < 0.001. Error bars represent SEM.

## Discussion

Although Zn^2+^ is known to play a critical role in mitochondrial homeostasis, it was not understood previously how mitochondrial Zn^2+^ is controlled at an appropriate level. In this study, we used unbiased genetic screens, combined with targeted gene screening, to reveal the identities of mitochondrial transporters that control mitochondrial Zn^2+^ homeostasis (Fig. 9 F). Our findings demonstrated that SLC-30A9 is the mitochondrial Zn^2+^ exporter. This is evidenced by the facts that SLC-30A9 localizes to mitochondria and binds with Zn^2+^, and its functional loss causes a strong elevation in mitochondrial Zn^2+^ levels and consequently severe damage of mitochondrial structure and functions. Loss of *slc-30A9* strongly impaired animal development, as revealed by the findings that *slc-30A9* mutants exhibit severely reduced fertility, embryonic lethality and retardation of larval development. In addition, Zn^2+^-induced mitochondrial damage greatly reduced the life span of *slc-30A9* mutants. These results suggest that SLC-30A9-mediated mitochondrial export of Zn^2+^ is critical for mitochondrial homeostasis, animal development and life span.

Mitochondrial calcium uniporter (MCU) was suggested to mediate mitochondrial entry of Zn^2+^ (Cho et al., 2019; Malaiyandi et al., 2005b; Medvedeva and Weiss, 2014). However, genetic inactivation of *mcu-1*, which encodes the *C. elegans* homolog of mammalian MCU, did not suppress the abnormal mitochondrial enlargement in *slc-30A9* mutants. This suggests that the mechanism of mitochondrial Zn^2+^ import is independent of MCU. Using two suppressor screen strategies, we revealed that mutations in *slc-25A25* suppressed the severe mitochondrial defects of *slc-30A9* mutants. Our results further indicated that both *C. elegans* and human SLC25A25 localize to mitochondria and bind with Zn^2+^. Moreover, SLC25A25 functions in an evolutionarily conserved manner, as heterologous expression of human SLC25A25 successfully substituted the function of *C. elegans* SLC-25A25 in double mutants of *slc-25A25* with *slc-30A9*. Loss of *slc-25A25* significantly reduced mitochondrial accumulation of Zn^2+^ in *slc-30A9* mutants. In addition, both *C. elegans* and human SLC25A25 bind to Ca^2+^. These findings suggest that SLC-25A25 is a Ca^2+^-binding regulator of mitochondrial Zn^2+^ import. Importantly, ectopic expression of *C. elegans* SLC-25A25 or human SLC25A25 caused enlargement of mitochondria with elevation of mitochondrial Zn^2+^ levels. This provides strong evidence that SLC25A25 promotes mitochondrial Zn^2+^ import, in addition to its originally reported function as a mitochondrial ATP-Mg^2+^/Pi transporter (del Arco and Satrustegui, 2004; Fiermonte et al., 2004; Monne et al., 2017; Satrustegui et al., 2007) . Altogether, these findings suggest that SLC-25A25 and SLC-30A9 are a pair of mitochondrial transporters which control mitochondrial Zn^2+^ import and export, respectively, to maintain the mitochondrial Zn^2+^ levels required for normal mitochondrial structure and functions (Fig. 9 F). It was reported that human and plant ATP-Mg/phosphate carriers (APCs), such as hAPC1/SLC25A24 and Arabidopsis APC3, co-transport divalent cations including Mg^2+^, Mn^2+^, Fe^2+^, Zn^2+^, and Cu^2+^ in proteoliposome assays (Monne et al., 2017). In our study, ectopic expressed human and *C. elegans* SLC25A25 greatly increased mitochondrial Zn^2+^ levels, suggesting that SLC25A25 is very likely to transport Zn^2+^, in addition to its function as an ATP-Mg transporter. Further studies will be needed to demonstrate whether SLC25A25 functions as a Zn^2+^ transporter or an ATP-Mg/phosphate transporter that also transports zinc. It also remains to be determined whether Zn^2+^ is transported in complex with ATP.

Using targeted gene screening, we found that inactivation of *slc-30A5*/ZnT5 strongly suppressed the enlargement of mitochondria in *slc-30A9* mutants. In both *C. elegans* and HeLa cells, worm SLC-30A5 and human SLC30A5/ZnT5 localize to the ER and bind Zn^2+^, suggesting that they function on the ER for Zn^2+^ transport. We found that RNAi of *slc-30A5*/*ZnT5* significantly reduced the Zn^2+^ levels in both ER and mitochondria in *C. elegans*. This suggests that SLC-30A5 functions as an ER Zn^2+^ importer and that the ER serves as the Zn^2+^ reservoir from which mitochondrial Zn^2+^ is imported. Supporting this conclusion, we found that siRNA knockdown of *ZnT5* significantly inhibited the abnormal mitochondrial enlargement and elevated mitochondrial Zn^2+^ levels in HeLa cells ectopically expressing *C. elegans* SLC-25A25 or human SLC25A25. Our findings suggest that *C. elegans* SLC-25A25 plays an important role in mitochondrial Zn^2+^ import, consistent with other studies that mammalian SLC25A25 localizes to the inner mitochondrial membrane (Satrustegui et al., 2007; Tewari et al., 2012). We reason that the transfer of Zn^2+^ from ER to mitochondria is achieved through ER-mitochondrion contacts (Fig. 9 F). It is possible that another ER transporter is responsible for exporting Zn^2+^ at the contacts, and the Zn^2+^, either alone or in complex with ATP, is then swiftly imported into mitochondria by SLC-25A25. Supporting this hypothesis, SLC-25A25 was found to be enriched at ER-mitochondrial contacts when *slc-30A9* is deficient (Fig. 9 D). Nevertheless, our unbiased genetic screen and targeted gene screen did not identify any ZnT-type Zn^2+^ exporters on the ER. In *Drosophila*, SLC25A25 was found to act downstream of the ER-localized Ca^2+^-permeable channel TRPP2 (transient receptor potential channel polycistin-2) to mediate TRPP2 signaling to regulate mitochondrial metabolism (Hofherr et al., 2018). In the case of Zn^2+^ transfer from the ER to mitochondria, further investigation is needed to determine whether TRPP2 or other transporters act upstream of SLC-25A25 to export Zn^2+^ out of the ER.

A single mutation in *SLC30A9* was recently found to be associated with an autosomal recessive cerebro-renal syndrome, which manifests as neurological deterioration, intellectual disability, ataxia, camptocormia, oculomotor apraxia, and nephropathy (Perez et al., 2017). However, the molecular and cellular basis for this genetic disorder remains unknown. Deregulation of *SLC30A9* expression was also reported for hepatocellular carcinoma (HCC) (Gartmann et al., 2018) and prostate cancer (PCa) (Singh et al., 2016). Likewise, *SLC25A25* expression was found to be mis-regulated in fulminant type I diabetes (Ye et al., 2020), HCCs (Kido and Lau, 2019) and hypospadias (Karabulut et al., 2013). Our findings that SLC-30A9 and SLC-25A25 cooperate to control mitochondrial Zn^2+^ levels thus provide important mechanistic insights for understanding these human diseases.

## Materials and Methods

### *C. elegans* strains and genetics

*C. elegans* strains were grown and maintained on NGM medium seeded with *E. coli* OP50 at 20°C. The Bristol strain N2 was used as the wild type. The Hawaiian strain CB4856 was used in single nucleotide polymorphism (SNP) mapping. *slc-30A9(yq158*), *slc-30A9(yq166*), *slc-30A9(yq172*), *slc-30A*(*yq189*), *slc-30A9(yq212*), *slc-30A9(yq158*);*slc-25A25(yq350*), *slc-30A9(yq158*);*slc-25A25(yq351)*, *slc-30A9 (yq158*);*slc-25A2 (yq371*), *slc-30A9(yq158*);*slc-25A25(yq372*) and *slc-30A9(yq158*); *slc-25A25(yq373*) were obtained by ethyl methanesulfonate (EMS) mutagenesis. *slc-25A25* (*yq406*) mutants were generated by the CRISPR/Cas9 system. *zcIs13*(P*_hsp-6_GFP*) was provided by the Caenorhabditis Genetics Center (University of Minnesota, Minneapolis, MN). The *mcu-1* (*ju1154*) deletion mutant were obtained from S. H. Xu (Center for Stem Cell and Regenerative Medicine, Zhejiang University School of Medicine, China).The integrated arrays *yqIs157* (P *_Y37A1B.5_mito-GFP*), *yqIs179* (P*_Y37A1B.5_tomm-20::mCherry*), and *yqIs180*(P*_mec-7_MTS::GFP*) were generated in this laboratory. Integrated arrays, deletion strains, and mutants generated by EMS mutagenesis or CRISPR/Cas9 were outcrossed with the N2 strain at least four times.

Extrachromosomal transgenes were generated by using standard microinjection methods. ≥3 independent transgenic lines were analyzed per construct. Transgenes used in this study are: *yqEx1196* (P*_F54A3.5_F54A3.5::GFP*), *yqEx1193* (P*_col-19_slc-30A9::GFP*), *yqEx1195* (P*_slc-30A9_slc-30A9*), *yqEx1296* (P*_slc-30A9_slc-30A9*(W305stop)), *yqEx1271* (P*_slc-30A9_gfp*), *yqEx1030* (P*_col-19_slc-30A9*(D198A;D202A)*::GFP*), *yqEx1342* (P*_col-19_slc-30A9*(D221A)*::GFP*), *yqEx1345* (P*_col-19_slc-30A9*(H402A)*::GFP*), *yqEx1349* (P*_Y37A1B.5_slc-25A25::mCherry*), *yqEx1352* (P*_Y37A1B.5_slc-25A25::GFP*), *yqEx1515* (P*_slc-25A25_slc-25A25::GFP*), *yqEx1516* (P*_slc-25A25_hSLC25A25::GFP*), *yqEx1517* (P*_tram-1_tram-1::BFP*), *yqEx1518* (P*_Y37A1B.5_*Mito*-*eCALWY-4), *yqEx1519* (P*_Y37A1B.5_*ER*-*eCALWY-4).

### Expression constructs

Bacterial, *C. elegans*, and mammalian expression constructs are listed in Table S1.

### RNA interference (RNAi)

RNAi experiments were performed by using the standard feeding method. For most experiments, 3-5 L4 larvae (P0) were placed on plates seeded with bacteria expressing gene-specific double-strand RNA and cultured at 20°C. The F1 progeny at the L4 stage were transferred to fresh RNAi plates or control RNAi plates. The adults after L4 (F1) 24-48 h were used for further analysis. For *slc-30A5* RNAi, 20 L4 larvae were placed on control (ctrl) RNAi and *slc-30A5* RNAi plates and cultured at 20°C. 24 h later, the adults were removed. Embryos were then analyzed for viability or developmental time.

### EMS screening and gene cloning

Synchronized L4-stage animals were treated with 50 mM EMS for 4 h. The F2 progeny at 24-48 h after the L4 molt were examined for phenotypes of interest. *yq158, 166, 172, 189* and *212* were isolated from a screen of about 10,000 haploid genomes. *yq158* was mapped to the genetic position from -12 to -10.69 on linkage group III (LGIII) using SNP mapping. The underlying mutations of all the mutants were identified with genome sequencing.

For the *slc-30A9*(*yq158*) suppressor screen, each of about 3,000 haploid genomes were screened for suppressors of mitochondrial enlargement and for suppressors of UPR^mt^ (P*_hsp-6_GFP*) signals. Chromosome mapping and gene identification were performed as above.

### CRISPR/Cas9-mediated gene editing

To generate *slc-25A25(yq406)* mutants, a single-guide RNA (sgRNA) targeting sequence (5’-CTGAATCTCCGTCGCTACG-3’) in the first exon of the *slc-25A25* gene was cloned into the pDD162 vector, which expresses the Cas9 enzyme. A repair template containing the mutation of interest was designed to remove the cleavage site. A restriction site (*Nhe I*) was introduced into the repair template. *dpy-10* was used as a positive control marker as described previously (Paix et al., 2014). pDD162 containing the sgRNA sequence (20 ng/μl) and repair templates (2 μM) for the target gene were coinjected with the *dyp-10* sgRNA construct (20 ng/µl) into gonads of young adult animals. Roller F1 worms were singled into new NGM plates, and the F2 progeny were examined by PCR amplification and restriction digestion. All mutations were confirmed by sequencing.

### Predication of protein structure

A structural model of the *C. elegans* SLC-30A9 protein (residues 152-495) was predicted using the Phyre2 protein fold recognition server (Kelley et al., 2015) (template PDB accession No. 3J1Z). Extremely conserved residues that could be confidently predicted to coordinate Zn^2+^ were labeled and shown as sticks. Zinc ions were shown as magenta spheres. The picture was prepared using PyMOL (The PyMOL Molecular Graphics System, version 1.8.0; Schrödinger).

### Cell culture, transfection, and reagents

HEK293 and HeLa cells were cultured at 37°C with 5% CO_2_ in DMEM (Gibco) with 10% heat-inactivated FBS (BioInd), 100 U/ml penicillin, and 100 µg/ml streptomycin. Transient transfections were performed with Lipofectamine 2000 (Invitrogen) according to the manufacturer’s instructions. MitoTracker Deep Red (Thermo) staining was used to indicate the location of mitochondria in live cells.

### Small RNA interference (siRNA)

Human *SLC30A5/ZnT5* siRNA oligos used in the study were siRNA1: 5’-GCUGGAGUAAACAAUUUAATT-3’, siRNA2: 5’-GGCUGAUAGUAAACCUUAUTT-3’, siRNA3: 5’-GUGGUAGUGAGUGCUAUAUTT-3’; Control siRNA: 5’-UUCUCCGAACGUGUCACGUTT-3’.

Mixed oligos of siRNA 1, 2 and 3 (100 pmol for each) or control siRNA were transfected into HeLa cells using Lipofectamine 2000 (Invitrogen). 48 h after transfection, cells were observed under fluorescence microscopy or harvested for qPCR to evaluate the knockdown of *SLC30A5/ZnT5*. qPCR oligos for examining *SLC30A5/ZnT5* expression were Forward: 5’-TTCTCCTATGGGTACGGCCGAAT-3’, Reverse: 5’-AGCCCTCCAACTGAGACTGGTGTT-3’.

### Microscopy and imaging analysis

Differential interference contrast (DIC) and fluorescence images were taken by an inverted confocal microscope (LSM880; Carl Zeiss) using a 100× 1.46-NA oil objective lens. Imaging was performed on animals of the same developmental stage and on a similar body region of each animal, unless otherwise indicated. Images were processed and analyzed with ZEN 2 blue software (Carl Zeiss) or Image J (National Institutes of Health). All images were taken at 20°C.

### MitoTracker Red CMXRos and Zinpyr-1 treatment

For MitoTracker Red CMXRos staining in *C. elegans*, the synchronized animals (24 h after L4 molt) were soaked in 50 µl MitoTracker Red CMXRos (Invitrogen, 10 µM in M9 buffer) for 10 min at 20°C in the dark. The worms were then transferred to a new OP50-seeded NGM plate for 2 h at 20°C in the dark and examined by fluorescence microscopy.

To visualize mitochondrial Zn^2+^ in HeLa cells, the cell-permeable fluorogenic Zn^2+^ reporter Zinpyr-1 (ChemCruz Biochemicals) was used by following the manufacturer’s instructions. Briefly, a 1 mM stock solution of Zinpyr-1 was diluted to a final concentration of 10 μM in calcium- and magnesium-free PBS buffer. Before treatment, cells were washed by calcium- and magnesium-free PBS buffer 3 times, and then incubated with the diluted Zinpyr-1 solution for 20 min at 37□°C with 5% CO_2_. After extensive washing, imaging was performed with fluorescence microscopy.

### ATP measurements

500 synchronized animals (24 h after L4 molt) were picked to a tube contain 100 µl lysis buffer (ATP-Lite Assay kit; Vigorous). The mixture was frozen in liquid nitrogen for 5 min and then heated at 100 °C for 15 minutes. The mixture was cleared at 5,000 rpm for 5 min at 4 °C and the supernatant was applied to the luciferin-luciferase assays in the ATP-Lite Assay kit.

### Life span assays

L4-stage worms were picked to fresh NGM plates and transferred to new plates every day until reproduction ceased. Worms were scored as dead if they had no response to touches on the head or tail. The surviving animals were recorded every day. Three repeats were performed for every strain.

### Bending frequency analysis

Worms at the same developmental stage were picked onto an uncoated NGM plate and allowed to crawl free from any adherent food. Worms were then transferred to another uncoated NGM plate and soaked in M9 buffer. The number of body bends generated in a 1 min time interval were scored by eye. A body bend was defined as a change in the direction along the longitudinal axis from the head to the tail. The animals were recorded individually with the investigator blind to the genetic status of the worms.

### Transmission electron microscopy

Adult animals were fixed using a high-pressure freezer (EM ICE, Leica). Freeze-substitution was carried out in anhydrous acetone containing 1% osmium tetroxide and 0.1% uranyl acetate dihydrate. The samples were sequentially placed at -90°C for 72 h, -60°C for 10 h and -30°C for 10 h, and 0°C for 5 h in a freeze-substitution unit EM AFS2 (Leica, Germany). After 3 washes (20 min each) with fresh cold anhydrous acetone, the samples were infiltrated with Embed-812 resin. Samples were embedded at 60°C for 48 h and cut into 70 nm sections with a microtome (EM UC7, Leica, Germany). After electron staining with uranyl acetate and lead citrate, sections were observed with a HT7800 TEM (HITACHI, Japan) operating at 80 kV.

### Recombinant proteins

cDNAs of interest were fused with the open reading frame of GFP and cloned into the pGEX-4T1 vector and transformed into the *E. coli* Rosetta (DE3) strain. The bacteria were first grown at 37°C to an OD_600_ of 0.6 and then added with 0.8 mM IPTG (isopropyl b-D-1-thiogalactopyranoside). The bacteria were then grown at 18°C for 24 h to allow for expression of soluble proteins. Recombinant proteins listed below were extracted in the extraction buffer containing 20 mM Tris-HCl, pH 8.0, 150 mM NaCl, 0.1% NP-40, 10% glycerol, 1 mM PMSF (phenylmethylsulfonyl fluoride) (Roche) and purified with Glutathione Sepharose 4B beads (GE Healthcare) according to the manufacturer’s instructions. GST-fusion proteins immobilized on the beads were eluted with Thrombin (Solarbio) in the protein buffer (25 mM HEPES, pH 7.4, 150 mM NaCl) for 6 h at 37 °C to obtain recombinant proteins without the GST tag. Recombinant proteins tagged with EGFP were used for the MST assays, including: the Control protein, EGFP. *C. elegans* protein, SLC-25A25-EGFP. Human protein, SLC25A25-EGFP.

### Isolation and purification of *C. elegans* mitochondria and Zn^2+^ measurement

Isolation and purification of *C. elegans* mitochondria were essentially performed as described previously (Li et al., 2009). Briefly, young adult worms obtained by liquid culture were collected and cleaned with 0.1 M NaCl, followed by centrifugation in 30% sucrose solution at 3,500 rpm for 5 min at 4°C. The upper layer containing the adult worms was transferred to fresh solution containing 0.1 M NaCl for further cleaning. Worms were then disrupted with ultra-sonication and subjected to differential centrifugation to obtain the crude mitochondria fraction. Mitochondria were purified with sucrose density gradient centrifugation using 55%, 40% and 30% sucrose gradients and 135,000 g at 4°C for 2 h. The mitochondrial layer was collected and further washed, and examined by western blotting using antibodies against mitochondrial, lysosomal, and ribosomal proteins (provided by Dr. Xiaochen Wang, Institute of Biophysics, Chinese Academy of Sciences).

Mitochondrial contents were released in RIPA buffer and analyzed with ICP-MS (Inductively Coupled Plasma Mass Spectrometry) in the Toxicology Department of Peking University School of Medicine and Tsinghua University.

### Microscale thermophoresis (MST) assays

To examine the binding of divalent cations with purified recombinant proteins, proteins at 200 nM were incubated with varying concentrations of CaCl_2_ and ZnSO_4_ in a 10 µl mixture and loaded into NT.115 standard coated capillaries (NanoTemper Technologies). The buffers (25 mM HEPES, pH 7.4, 150 mM NaCl) used in the MST assays were free from EDTA/EGTA. MST measurements were performed at 25 °C, 60 % excitation power, and medium MST power. All experiments were repeated three times for each measurement. Data analyses were performed using NanoTemper software.

To examine cation binding with EGFP-fused proteins expressed in HEK293 cells, 10 µl of cell lysates containing the protein of interest in lysis buffer (25 mM Tris-HCl, pH 7.5, 100 mM NaCl, 1% NP-40, 1% glycerol, 1 mM protease inhibitor cocktail-EGTA free, and 1 mM PMSF) were incubated with varying concentrations of divalent cations (ZnSO_4_, CaCl_2_, MgSO_4_, CuSO_4_, MnCl_2_) and loaded into NT.115 standard coated capillaries. MST measurements and analysis were performed as above.

### Fluorescence resonance energy transfer (FRET) imaging

The *C. elegans* Mito-eCALWY-4 construct was made by fusing 4 repeats of the mitochondrial targeting sequence of human cytochrome C oxidase subunit VIII (Cox VIII) to the N-terminus of eCALWY-4 and cloning into the pPD49.26 vector (Fig. 2 G). The *C. elegans* ER-eCALWY-4 construct was generated by fusing the preproinsulin (PPI) sequence and the Lys-Asp-Glu-Leu (KDEL) sequence at the N- and C-terminus of eCALWY-4, respectively (Fig. 2 G). Expression of both probes was controlled by the *C. elegans* hypodermal *Y37A1B.5* promoter. Mito- and ER-eCALWY-4-expressing constructs were microinjected into N2 animals to obtain the extrachromosomal transgenic lines *yqEx1518*(P*_Y37A1B.5_*Mito*-*eCALWY-4) and *yqEx1519*(P*_Y37A1B.5_*ER*-*eCALWY-4). All strains expressing Mito-eCALWY-4 or ER-eCALWY-4 probes were obtained by introducing these extrachromosomal arrays.

Synchronized animals (24 h after L4 molt) were imaged using a confocal laser scanning microscope (LSM880; Carl Zeiss) with a 100× 1.46-NA oil objective lens. Cerulean (CFP) and Citrine (YFP) were excited at 458 and 514 nm, respectively, and the fluorescence emission was detected at 450-490 and 520-560 nm, respectively. The FRET channel was excited at 458 nm and fluorescence emission was detected at 520-560 nm. Normalized measure of FRET (FRETN) was performed as previously described (Gordon et al., 1998).

### Fluorescence lifetime imaging microscopy (FLIM)-FRET

FLIM-FRET imaging for eCALWY-4 probes was performed on a Leica TCS SP8X scanning confocal microscope equipped with a 63×/1.3-NA water immersion objective, a white light laser (WLL), a HyD SMD single molecule detector and a 405- nm pulsed-laser excitation source. The cerulean donor (CFP) was excited by a 405-nm picosecond pulsed diode laser tuned at 40 MHz. Emitted photons passing through the 460-500-nm emission filter were detected using the HyD detector. The pixel frame size was set to 512×512, speed for 200, fluorescence lifetime frame repetition was acquired over 20, and the maximal photon counting rate was around 1×10^6^ counts. The fluorescence lifetime was analyzed using the Leica FALCON FLIM integrated software. “τ_c_” indicates the fluorescence lifetime of the CFP donor.

### Statistical analysis

The two-tailed unpaired Student’s *t* test was used to ascertain statistically significant differences between two groups. One-way ANOVA with a Newman-Keuls post-test was used to ascertain statistically significant differences between multiple groups. For all quantifications, *, *P* < 0.05; **, *P* < 0.01; ***, *P* < 0.001; NS, not significant. Data were analyzed with GraphPad Prism Software (8.0.1) to generate curves or bar graphics with SEM (standard error of the mean) as y-axis error bars. Data distribution was assumed to be normal but this was not formally tested.

## Online supplemental materials

Fig. S1 characterizes mitochondrial defects in *slc-30A9* mutants. Fig. S2 compares amino acid sequences between bacterial Yiip, *C. elegans* SLC-30A9 and human SLC30A9. Fig. S3 characterizes SLC-30A9**/**SLC30A9 localization and binding with divalent cations. Fig. S4 characterizes *C. elegans* SLC-25A25. Table S1 summarizes the expression constructs. Table S2 summarizes the RNAi-based screen for candidate Zn^2+^ transporters.

## Acknowledgements

This work was supported by grants from the National Science Foundation of China (91954204 and 31730053), the National Basic Research Program of China (2017YFA0503403), and Yunnan Province Science and Technology Department (#202001BB050077). C. Yang is supported by Program of Yunnan Province Leading Talents in Science and Technology. We thank Dr. I. Hanson for proofreading the manuscript; Dr. G. Rutter (Imperial College of London) for providing the eCALWY-4 constructs; D. X. Wang (IBP, CAS) for providing *C. elegans* protein antibodies; Dr. L. Chen (Tsinghua Univ.) for ICP-MS analysis of mitochondrial divalent ions HeLa cells; Dr. X. Ma (Zheng Zhou Univ.) for help with SLC-30A9 structure prediction; and Dr. S. Xu (Zhejiang Univ.) as well as the *Caenorhabditis* Genetic Center (CGC) for *C. elegans* strains.

## Author Contributions

T. Ma, L. Zhao, J. Zhang, and R. Tang performed most of the experiments and interpreted the data. X. Wang and M. Li performed TEM analysis. N. Liu, F. Wang, Q. Zhang, Q. Shan, Q. Yin, Y. Yang and Q. Gan contributed to the experiments and materials. T. Ma, J. Zhang and C. Yang prepared the manuscript with discussion between all authors. C. Yang designed and supervised the study.

## Declaration of interests

The authors declare no competing financial interests.

## Notes

### Competing Interest Statement

The authors have declared no competing interest.

