## Supplemental materials for "A pair of transporters controls mitochondrial Zn^2+^ levels to maintain mitochondrial homeostasis"

**Figure S1. Characterization of the mitochondrial defects in *slc-30A9(yq158)* mutants.**

(**A**) Images of mitochondria labeled with F54A3.5::GFP and TOMM-20::mCh in the hypodermis of N2 and *slc-30A9*(*yq158*) animals. Bars, 5 μm.

(**B**) Representative TEM images of mitochondria in the germline, sperm, oocyte and embryo of N2 and *slc-30A9*(*yq158*) animals. Bars, 1 μm.

(**C**) Time-course comparison of Mito-GFP-labeled mitochondria in the hypodermis of N2 and *slc-30A9*(*yq158*) animals. Bars, 5 μm.

(**D**) Expression of P*_slc-30A9_slc-30A9*::*GFP* in hypodermal, muscle and intestinal cells as well as in the PVD neuron in N2 adults. DIC and ﬂuorescence images are shown for each tissue. Bars, 5 µm.

**Figure S2.** Amino acid sequence alignment of bacterial (*E.c*) Yiip, *C. elegans* SLC-30A9 and human (*H.s*) SLC30A9. Residues important for Zn^2+^ binding are indicated in red boxes. Mitochondrion-targeting sequences (MTSs) are indicated in blue boxes. The cation-efflux domains are indicated in pink boxes.

**Figure S3. Characterization of SLC30A9 localization and binding with divalent cations.**

(**A**) Localization of *C. elegans* SLC-30A9-EGFP, SLC-30A9(1-145)-EGFP, and SLC-30A9(146-495)-EGFP in HeLa cells. Mitochondria are labeled with MitoTracker DR. Boxed regions are magnified (2.5×) in the insets. Bars, 2 µm.

(**B**) Binding curves of HEK293 cell-expressed SLC-30A9-EGFP with divalent cations (Ca^2+^, Mg^2+^, Cu^2+^ and Mn^2+^), measured with MST assays. EGFP was used as the negative control.

(**C**) Co-localization of human SLC30A9-EGFP with MitoTracker DR in HeLa cells. Boxed regions are magnified (2×) in the insets. Bars, 2 µm.

(**D**) Binding curve of HEK293 cell-expressed human SLC30A9-EGFP with Zn^2+^, measured with MST assays. EGFP was used as the negative control.

**Figure S4. Characterization of *C. elegans* SLC-25A25.**

**(A)** Amino acid sequence alignment of *C. elegans* (*C. e*) SLC-25A25 and human (*H. s*) SLC25A25. EF-hands are indicated in red boxes. Mitochondrial carrier domains are indicated in blue boxes.

**(B)** Expression pattern of P*_slc-25A25_slc-25A25*::*GFP* in *C. elegans.* DIC and ﬂuorescence images are shown for the indicated tissues. Bars, 5 µm.
