## Supplementary figures and images for "A pair of transporters controls mitochondrial Zn^2+^ levels to maintain mitochondrial homeostasis"

### Supplemental Figures

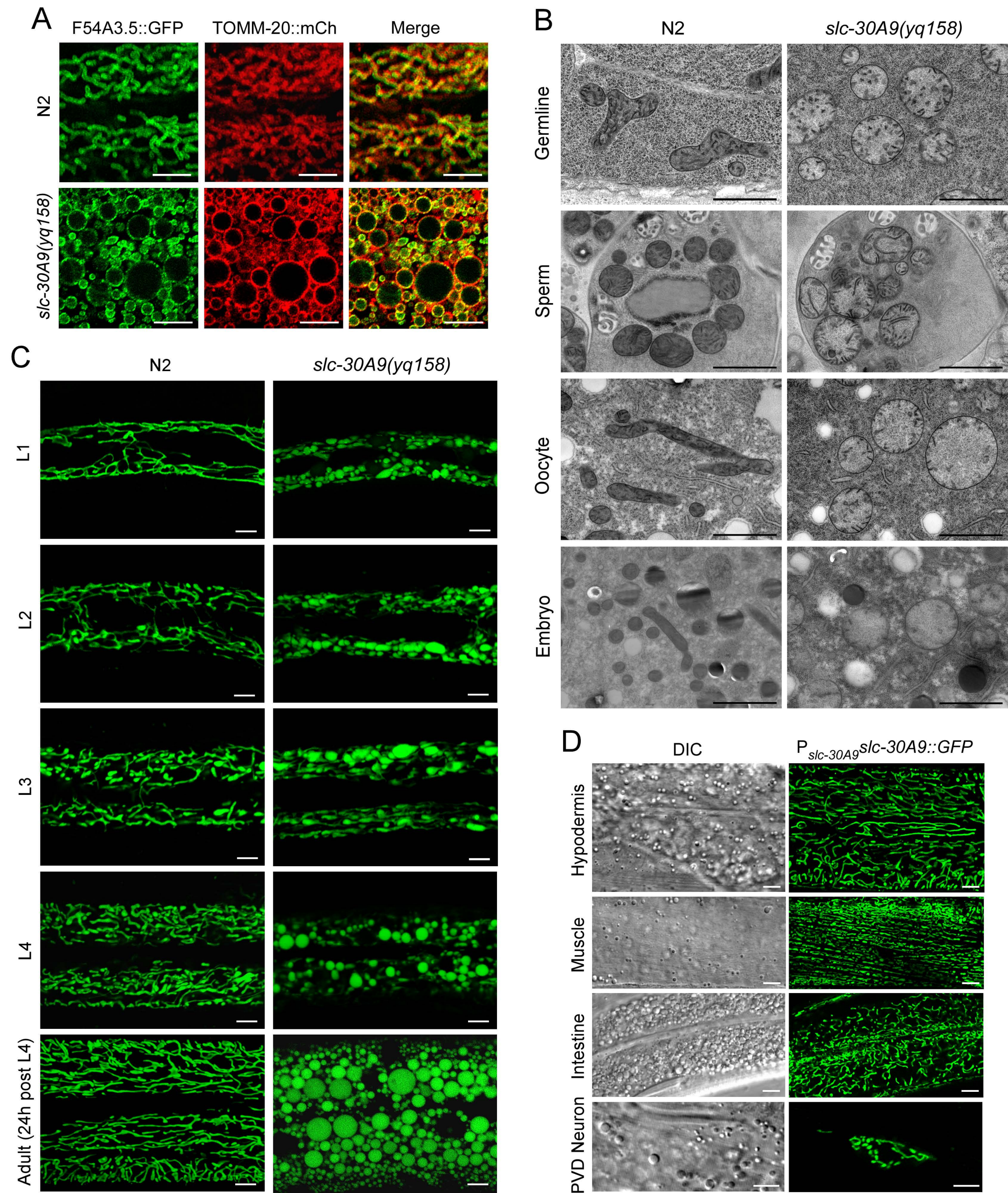

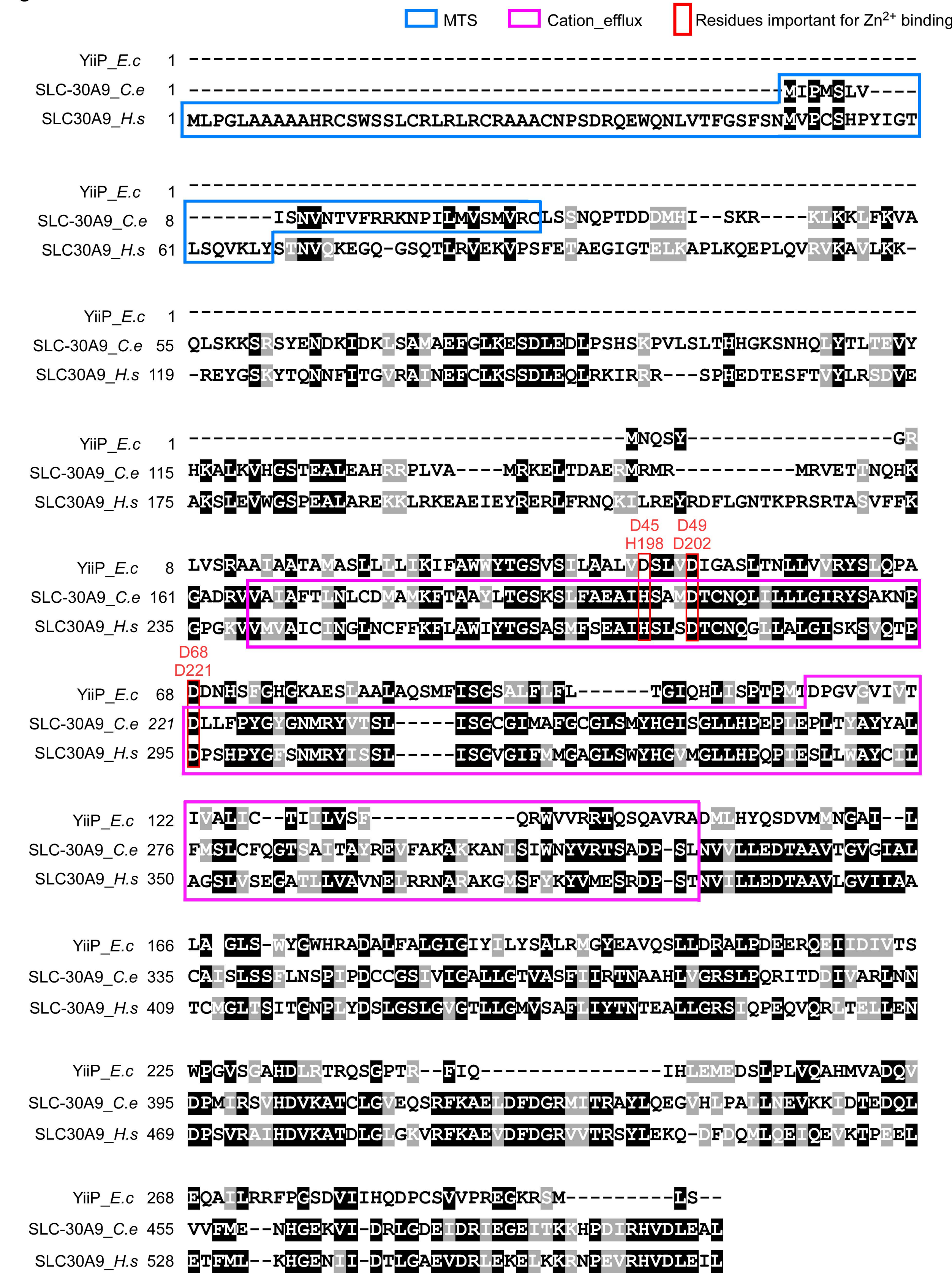

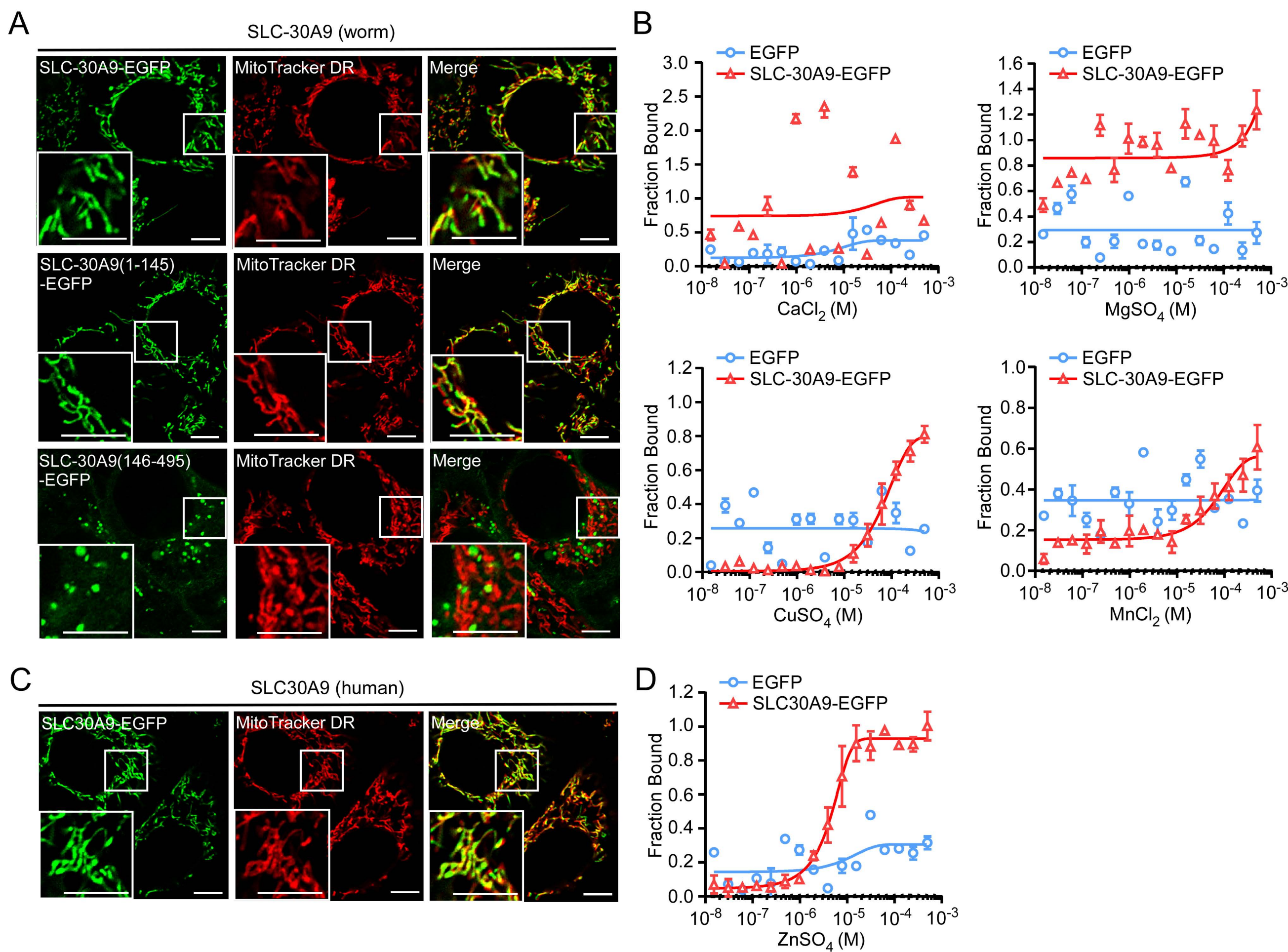

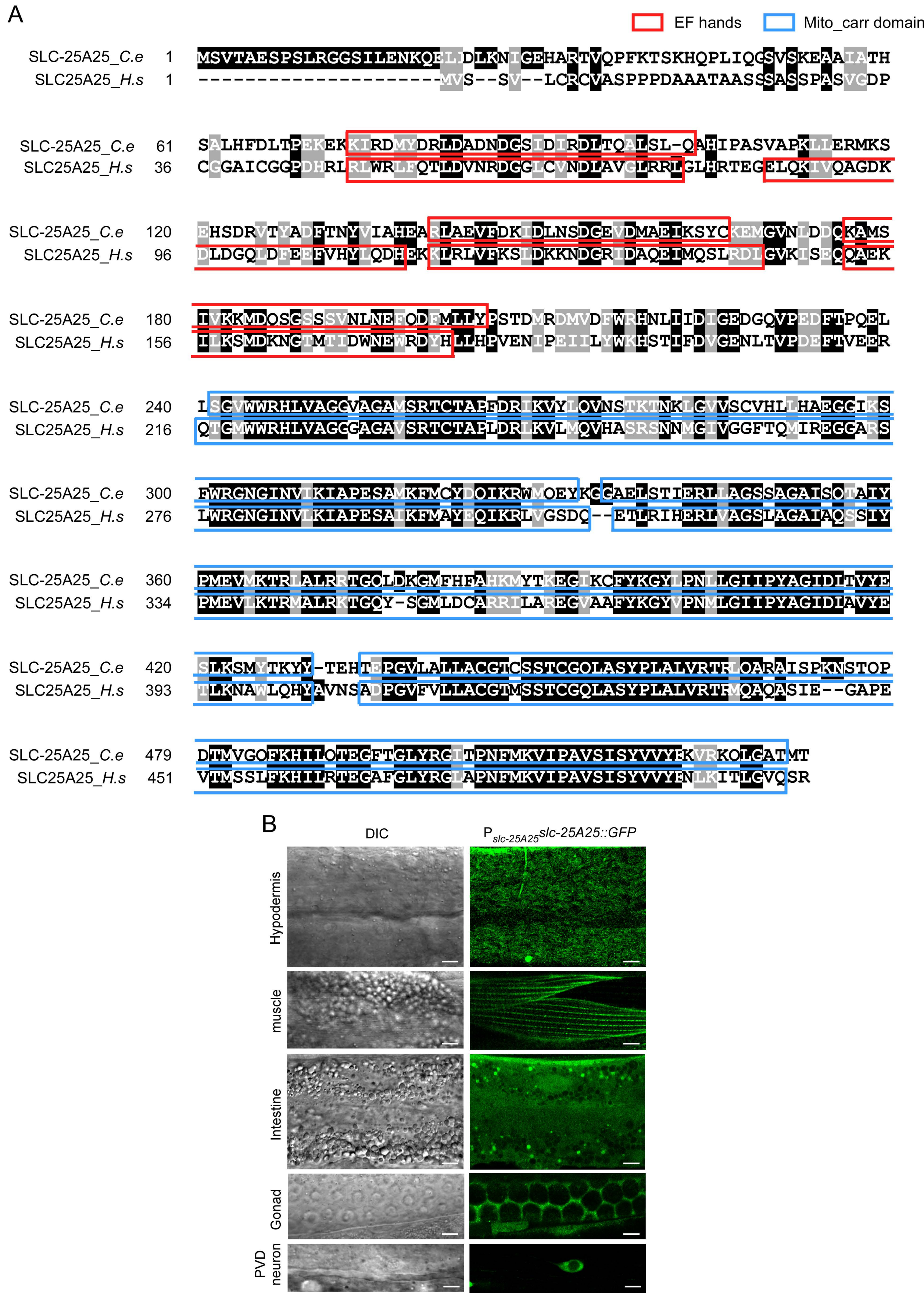
